# Quantifying invasibility

**DOI:** 10.1101/2021.06.22.449376

**Authors:** Jayant Pande, Yehonatan Tsubery, Nadav M. Shnerb

## Abstract

Invasibility, the chance of a population to grow from rarity and to establish a large-abundance colony, plays a fundamental role in population genetics, ecology, and evolution. For many decades, the mean growth rate when rare has been employed as an invasion criterion. Recent analyses have shown that this criterion fails as a quantitative metric for invasibility, with its magnitude sometimes even increasing while the invasibility decreases. Here we employ a new large-deviations (Wentzel-Kramers-Brillouin, WKB) approach and derive a novel and easy-to-use formula for the chance of invasion in terms of the mean growth rate and its variance. We also explain how to extract the required parameters from abundance time series. The efficacy of the formula, including its accompanying data analysis technique, is demonstrated using synthetic and empirically-calibrated time series from a few canonical models.

---

The capacity of natural systems to maintain biodiversity (for example, genetic polymorphism and coexistence of multiple competing species) poses a major challenge to the theory of population genetics and community ecology [1, 2]. Interspecific interactions in diverse communities reflect many complex factors, and their effects are intertwined with strong fluctuations caused by environmental and demographic stochasticity. A reliable inference of model parameters, which is required for evidence-based analysis, is therefore a formidable task.

Turelli’s [3, 4] approach to this problem focuses on mutual invasibility, i.e., on the “conditions under which a rare invading species will tend to increase when faced with an array of resident competitors in a fluctuating environment”. This shift in focus greatly simplifies the study, as the complicated system reduces to a series of single-species invasion problems. All complex interactions are then encapsulated in a few effective parameters that reflect the overall influence of the resident species and the environmental fluctuations on a given rare population. The rarity of the focal species allows one to neglect nonlinear (density-dependent) effects, which facilitates the analysis even further. An analogous approach is taken in permanence (uniform persistence) theories [5–8].

Chesson’s modern coexistence theory [9] bases its invasibility analysis on a *single* parameter, the mean growth rate of the invader species 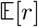 [3, 10, 11]. The mean is taken over all the instantaneous growth rates *r*(*t*) where *t* denotes time. The same parameter, 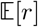, has been employed in the adaptive dynamics theory of evolution [12, 13] and in many other fields [14]. Consequently, ecologists have developed a collection of techniques to infer 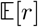 from empirical time series or from empirically calibrated models [14]. Many contemporary studies of species coexistence and the maintenance of biodiversity analyze this “invasion criterion” 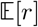 [14] and partition it between underlying mechanisms like niche differentiation, fitness differences, effects of the fluctuating environment and so on [15–17]. In parallel, many authors employ 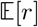-based criteria for mutual invasibility as a metric for stability in empirical studies [18, 19].

However, recent works have revealed that 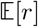 is not a reliable *quantitative* indicator of invasibility [20, 21]. Systems with different underlying parameters may yield different probabilities of invasion for a species despite having the same numerical value of 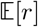. Even worse, systems may exhibit a decreasing invasibility even as 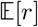 increases. Although 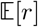 provides a fair *binary* classification – when its value is positive (negative), the extinction state is a repeller (attractor), from which important asymptotic predictions follow [6, 8, 11, 22, 23] – it does not reliably *measure* invasibility.

The failure of 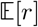 as a metric reflects two distinct problems [20, 23, 24]. First, since the growth rate *r* measures the logarithm of abundance ratios, zero-abundance (extinction) states lead to infinite negative contributions. Therefore, 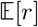-based analyses typically neglect demographic stochasticity (genetic drift), the intrinsic stochasticity arising from the birth and death of discrete individuals [25], despite its crucial importance when the invading population is small. Second, 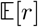 does not take fully into account the effect of environmentally-induced abundance variations – in other words, environmental stochasticity [25] – which affect entire populations coherently.

Thus, it is an open question to identify a quantitative metric that accurately predicts the chance of invasion of a rare species. Following former studies [25], Dean and Shnerb [21] suggested such a parameter for systems with environmental (but without demographic) stochasticity when the environmental fluctuations are weak enough for the diffusion approximation [26, 27] to be valid. However, the range of applicability of the diffusion approximation in varying environments (even when it takes demographic fluctuations into account) is rather narrow [28].

Here we provide a new and simple formula that predicts the chance of invasion. This formula works for weak as well as strong levels of both demographic and environmental stochasticity. Our formula converges to the known classical results for establishment probability [29–31] when the environment is fixed through time, and to the expression suggested in Ref. [21] when the diffusion approximation holds. As a bonus, our analysis clarifies how the effect of demographic stochasticity may be taken into account indirectly by introducing an “extinction threshold” at the right density. This allows one to extract quantitative predictions from infinite population models, like those used in permanence (uniform persistence) theories [5–8].

The derivation of our formula is based on a technique we developed [28] to extend the diffusion approximation by employing the WKB (Wentzel-Kramers-Brillouin) approximation scheme, building upon previous studies [32–35]. In general, the results of such an analysis are case-specific, since it must take into account nonlinear, density-dependent effects that manifest themselves in deterministic and stochastic parameters. In contrast, the invasion problem considered here is **universal**. During the invasion process the invader population is rare (its frequency is negligible when the size of the community is large) so all external parameters may be assumed to be fixed (density-independent) over the invasion regime. This simplification allows us to consider *all* possible dynamics and to present a universal formula that depends on four quantities: 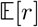 itself, the strength *V*_e_ and the typical duration *δ* of environmentally-induced abundance fluctuations, and the strength of demographic stochasticity *V*_d_. All these quantities may be extracted from a single species history, and the effort needed to calculate them does not differ significantly from that needed to calculate 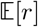 alone [as described in [14] and [15]].

The calculation procedure is illustrated in Figure 1. Consecutive censuses or long simulations, like those considered in [14, 15, 18], yield a time series {*n*_*t*_, *n*_*t*+*τ*_, *n*_*t*+2*τ*_, *n*_*t*+3*τ*_, …}, where *n* denotes the abundance of an invading species, *t* denotes the time (measured in units of a generation), and *τ* is the time interval between samplings. This dataset is translated to the log variable *z_t_* = ln *n_t_* [so *n_t_* = exp(*z_t_*)]. The dwell time *δ* is twice the autocorrelation time extracted from the *z* time series.

**FIG. 1:**
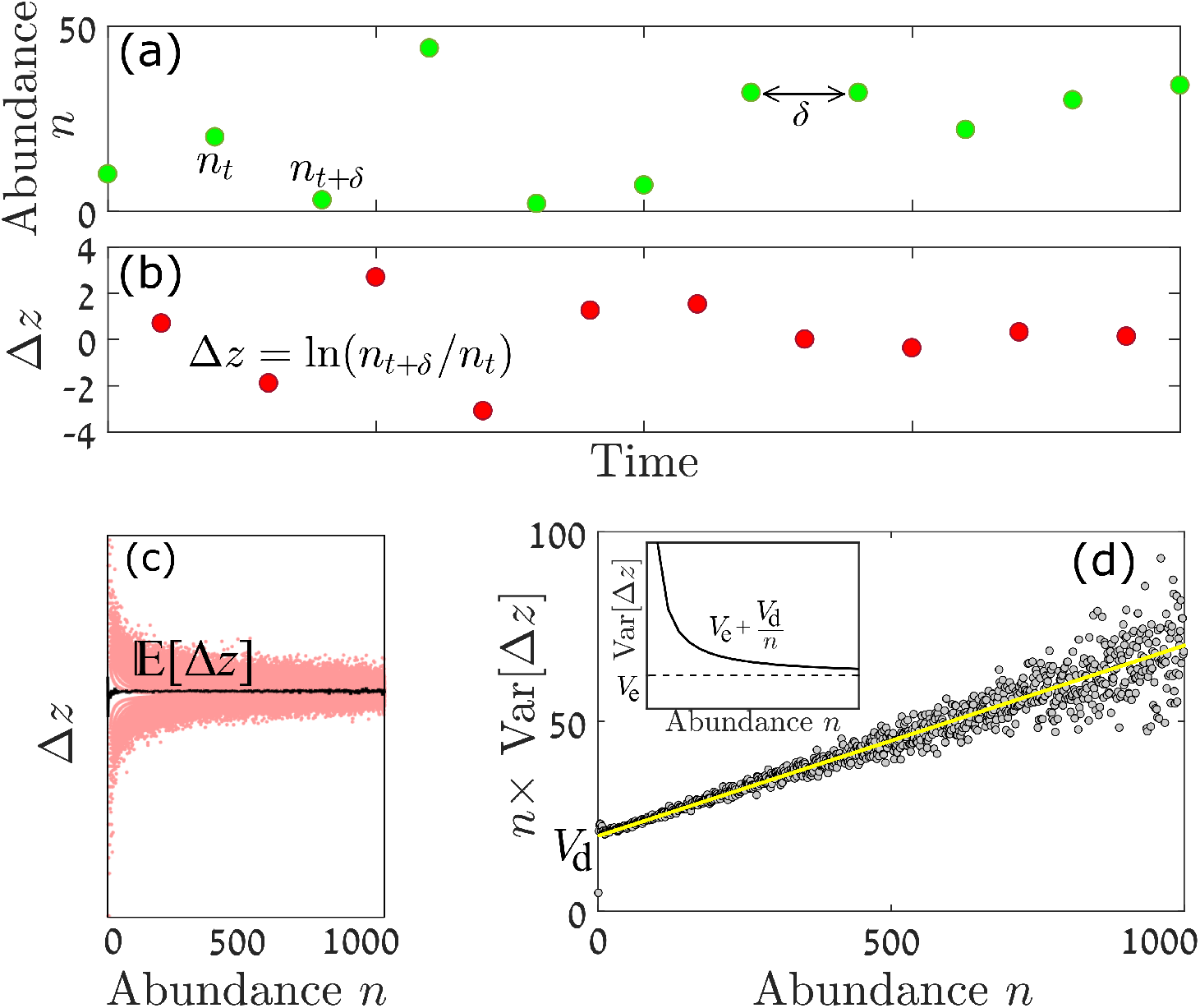
Calculating growth rate and stochasticity parameters. The raw data is a rarified abundance time series (as obtained from direct observations or a numerical simulation of an adequately calibrated model). Here this dataset of *n*_*t*_, *n*_*t*+*δ*_, *n*_*t*+2*δ*_, … is illustrated in panel (a). Each pair of points yields a single ∆*z* ≡ ln (*n*_*t*+*δ*_/*n*_*t*_) in panel (b). A scatter plot of ∆*z* values vs. *n* is presented in panel (c): the mean value of this quantity is, by definition, 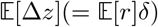 (black line). The scatter width decreases with *n*, since the demographic stochasticity component in the variance of ∆*z* decays like *V*_d_/*n*. The variance saturates to the level attributed to the environmental stochasticity, *V*_e_, when *n* is large [inset, panel (d)]. Multiplying the simulated variance by *n* and plotting it against the abundance yields a straight line whose intercept with the *y*-axis is *V*_d_ and whose slope is *V*_e_ [main part, panel (d)]. The datapoints shown in panels (c) and (d) were obtained from a numerical simulation of the discrete-time lottery model (see Methods) with *δ* = 0.1, *σ* = 0.2 and *s*_0_ = 0.1. The noisy behavior in the vicinity of the zero-abundance point has to do with extinction events that were not included in calculating the mean growth rate and its variance since they lead to divergence of ∆*z*.

Given *δ*, the time series is rarified [see Fig 1(a)] to obtain a series of ∆*z* changes over *δ*. The mean and the variance of this series yield 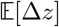, *V*_d_ and *V*_e_, via

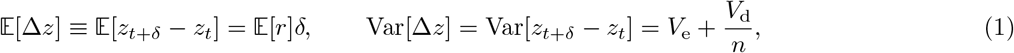

where Var denotes the variance. We distinguish between the abundance-independent part of this variance, *V*_e_, which reflects the environmental stochasticity, and the abundance-dependent part *V*_d_/*n*, which is due to demographic stochasticity and decays like 1/*n*. A partitioning of the variance, as explained in Figure 1, determines *V*_e_ and *V*_d_.

Once these quantities are found, the chance of invasion Π_*n*_ is given by

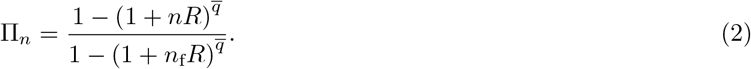

In this formula Π_*n*_ is the chance of a population of size *n* to grow in abundance to some prescribed value *n*_f_ before going extinct. The quantity 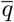 is given by

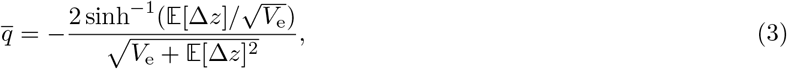

and *R* is defined as

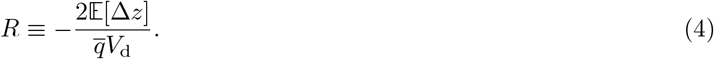

Eq. (2) is our central result. Its derivation is based on a technique presented in [28] and is explained in the Methods section and in section A in the Supplementary Materials. It assumes that in the invasion region (1 < *n* < *n*_f_), 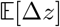, *V*_d_ and *V*_e_ are *n*-independent, which is generally true when *n*_f_ is a small fraction of *N*, the total population of the entire community counting all the species. When this assumption does not hold it is straightforward to replace Eq. (2) with an equally well-working, if less elegant, expression, involving an integration over the abundance *n* (Eq. (15)).

We demonstrate the effectiveness of our formula using simulated data from three individual-based versions of canonical models: the discrete-time lottery model of Chesson and Warner [36], its continuous-time (Moran) analog, and the Leslie-Gower competition model for trees and saplings as employed by [18, 19]. A short description of these models is provided in Methods. In all these models, in some ranges of parameters stochasticity promotes the coexistence of species (i.e., their mutual invasibility), a phenomenon often called the storage effect [2, 36]. This phenomenon makes the relationship between the underlying process parameters and the invasibility properties rather intricate and presents a demanding test to Eq. (2).

The efficacy of Eq. (2), and its superiority with respect to the use of 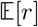 as a metric for invasibility, is demonstrated in Figure 2. Π_*n*=1_ in a discrete-time, individual-based version of the lottery model (see Methods) is plotted for different sets of values of the dwell time *δ* and the amplitude of environmental stochasticity *σ*. The left panel demonstrates the inefficacy of 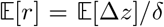 as an indicator, while the right panel shows how our formula predicts not only the general trend but also the numerical value of the chance of invasion to high accuracy.

**FIG. 2:**
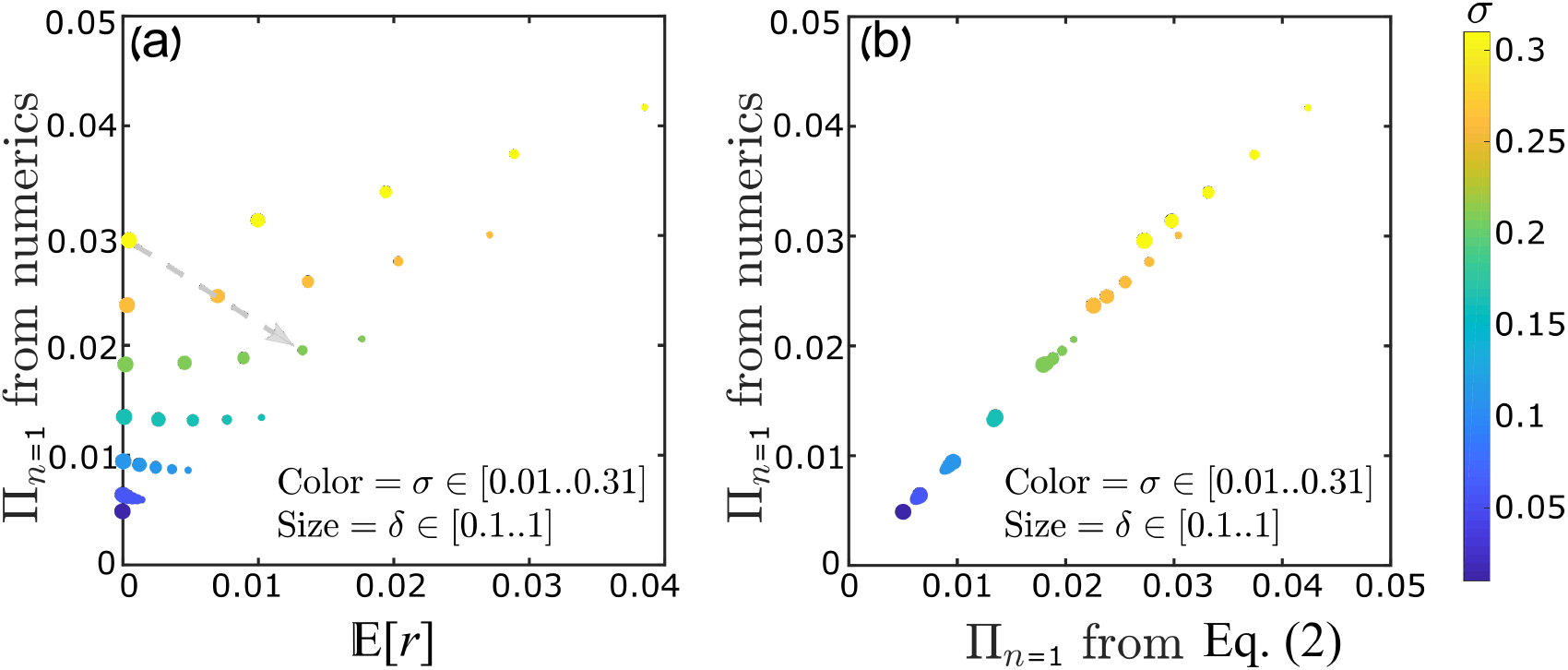
The chance of a single invader (*n* = 1) to reach *n*_f_ = 200 before extinction. This chance was obtained from numerical solution of the discrete-time lottery model with demographic stochasticity (see Methods) for various values of the model parameters *δ* (the dwell time) and *σ* (the amplitude of the temporal fitness variations). The color of each filled circle indicates the value of *σ*, as marked on the color scale shown, while its size is proportional to its *δ*-value. In the left panel the chance is plotted against 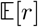, demonstrating that 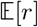 is a poor metric: by changing the model parameters *δ* and *σ* (for example, along the path indicated by the gray arrow) invasibility may decrease while 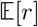 increases. In the right panel the same results are plotted against the predictions of Eq. (2), showing not only a data collapse but also close agreement with the numerical values. All the results were obtained for *s*_0_ = 0. Similar graphs for *n*_f_ = 20 and for *n*_f_ = 500 are presented and discussed in section C of Supplementary Materials.

Figure 3 shows Π_*n*_ as a function of *n* for the continuous-time (Moran) version of the lottery model for various values of *s*_0_ (the mean fitness advantage or disadvantage of the invading species). The analytical predictions are borne out by the numerical results over a wide range of values. In Figure 4 we present the comparison for the trees-saplings Leslie-Gower model employed by Usinowicz et al. [18], using their empirically-calibrated recruitment data (scaled by a constant factor) for the species *Spondias mombin* and *Spondias radlkoferi*. In the Leslie-Gower model, the tree dynamics involve time delays since seedling production in a given year affects the recruitment of trees for many years in the future. Therefore, while for the two versions of the lottery dynamics *δ* is a model parameter and 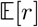, *V*_d_ and *V*_e_ are obtained analytically (as explained in section E of Supplementary Materials), in the Leslie-Gower case all these parameters must be calculated through the numerical steps illustrated in Figure 1 (section F of Supplementary Materials).

**FIG. 3:**
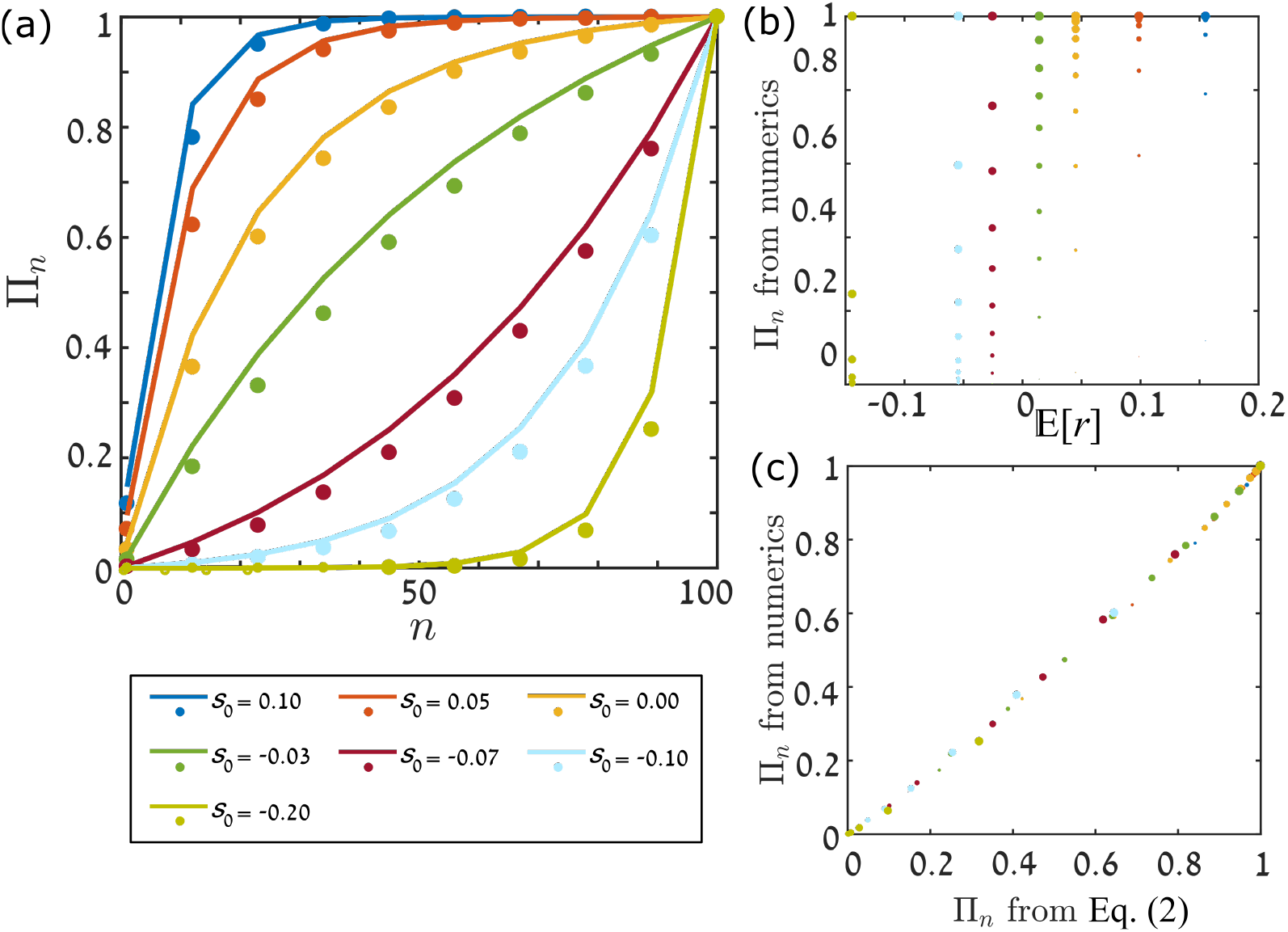
For the continuous-time (Moran) version of the lottery model, the chance to reach *n*_f_ = 200 starting from varying *n*, as obtained from Monte Carlo simulations (circles), is compared, in panel (a), with the predictions of Eq. (2) (full lines) for different values of *s*_0_, the mean selection parameter. In panel (b) the chance of invasion found from the simulations is plotted against 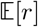, with the color of the points corresponding to their value of *s*_0_, and their size to the value of *n*. In panel (c) the same chance of invasion is plotted against the prediction of Eq. (2). As in Fig. 2, all the observed (in simulations) data points collapse on a line when plotted against Eq. (2). Other parameters are *σ* = 0.3, *δ* = 0.1 and *N* = 5000. Similar graphs for different values of *N* and *n*_f_ are presented in section D of Supplementary Materials. Even when *s*_0_ is negative, the curves in panel (a) may have the concave shape associated with a beneficial mutant. This happens because the storage effect [21, 36] may support invasion by an inferior species.

**FIG. 4:**
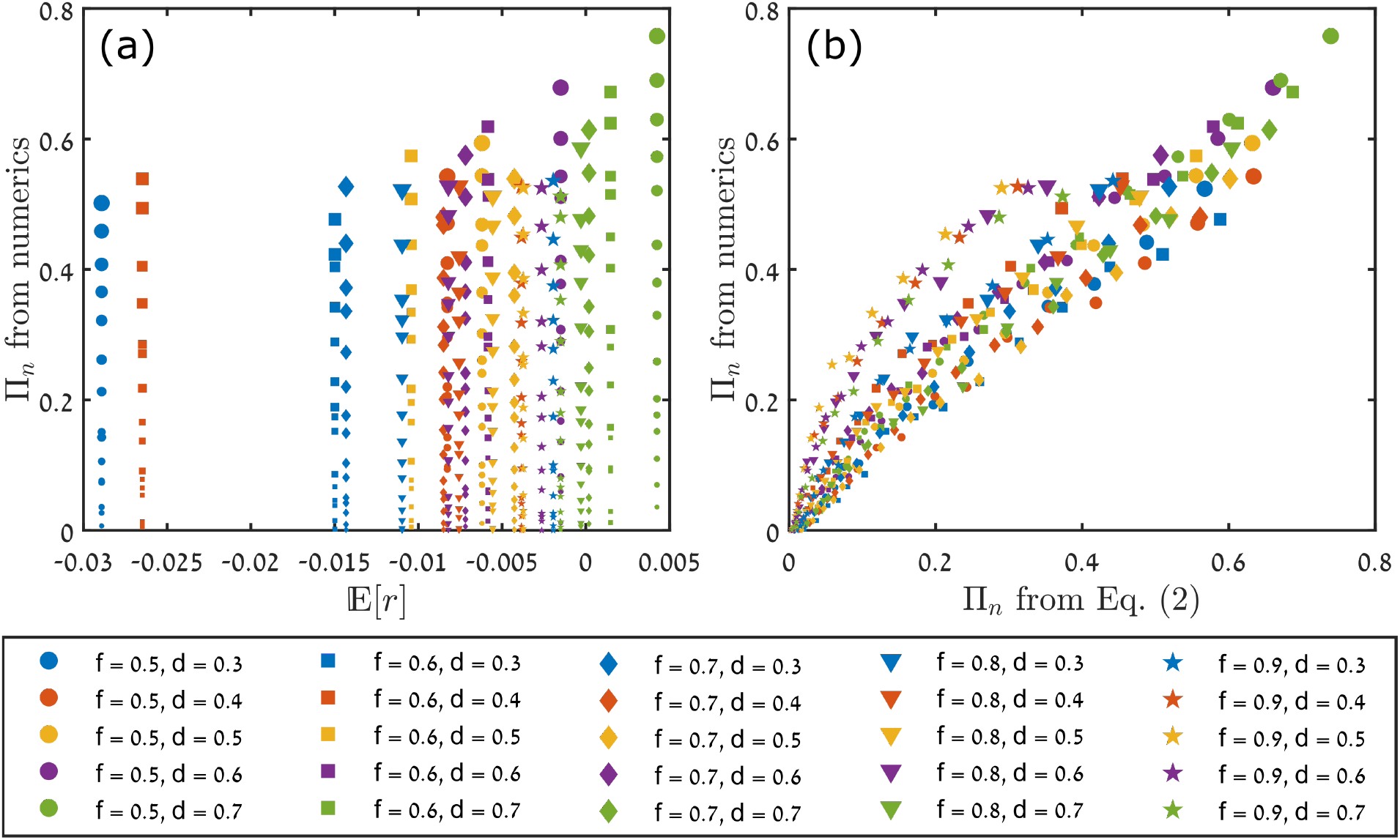
Chance of invasion in the Leslie-Gower forest dynamics model of [18, 19], as described in the Methods section. For a community size of *N* = 10000 we calculated the chance to reach *n*_f_ = 1000, starting from varying *n*, and plotted the results against 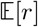 (panel a) and against the predictions of Eq. (2) (panel b). The size of each point is proportional to the initial population *n* ∈ [1‥200], different colors correspond to different chances of adult tree survival *d* ∈ [0.3‥0.7] and different symbols represent different chances of sapling survival *f* ∈ [0.5‥0.9]. In our simulations we employed the empirical recruitment rates for *Spondias mombin* and *Spondias radlkoferi*, as provided in Ref. [18], scaled by a constant factor. A step-by-step description of the analysis is presented in section F of Supplementary Materials.

Given the simplicity and accuracy of our parameter Π, we suggest that it replace 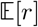 as an invasion metric everywhere in ecological and evolutionary analysis. As explained earlier, the additional parameters (namely *V*_d_ and *V*_e_) required for the calculation of Π can be inferred from the same datasets from which 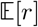 is calculated, and Π predicts the true chance of invasion much more accurately than 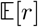. Our formula allows a quantitative estimation of the invasibility even when a direct numerical assessment (such as through simulations) is impossible or very difficult, e.g., when *n*_f_ is very large or when the chance of invasion is very close to 0 or to 1. This latter case is the situation in [19], where a comparison between the stability of different tree communities requires one to quantify the invasibility even though it is very close to 1.

When the environment is fixed in time (i.e. *V*_e_ = 0), Eq. (2) converges to known results that were obtained a long while ago by employing branching process theory [29] or the diffusion approximation [30], as explained in section B in Supplementary Materials.

Besides its success as a quantitative metric, our formula provides a few important and useful insights. First, Eq. (2) reveals the importance of the initial population *n* and the target abundance *n*_f_, two quantities that are rarely discussed in the literature. For example, when the initial abundance of the invader species is large, such that 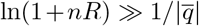, then the chance of invasion approaches unity when 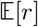 is positive [i.e., when 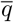 is negative, since 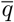 and 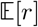 are of opposite signs, from Eq. (3)]. Under the same conditions the chance of invasion is 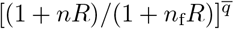 when 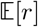 is negative (meaning 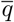 is positive). This means that the only case in which 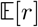 provides *quantitative* information about the chance of invasion is when 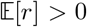 and the initial population is very large. In all other cases, and in particular when the invading population is small (as in the case of rare mutations, speciation events and epidemic outbreaks), the relevant analyses, like a mechanism-based partitioning of the chance of invasion, must be carried out with respect to Π.

A second insight provided by Eq. (2) is an informed definition of *n*_f_, the abundance level that defines a successful invasion. Two simple alternatives are choosing *n*_f_ to be some fixed large number (independent of the overall size of the community) or a fixed frequency (a given fraction of the community), but these choices are somewhat arbitrary. Our formula suggests that a natural definition for *n*_f_ (when 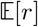 is positive so 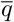 is negative) is the abundance above which the chance of extinction is small, 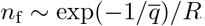.

Incorporating both demographic and environmental stochasticity in a model is a complicated task. Because of this, many population models, like those used in permanence (uniform persistence) theories [5–8], deal only with environmental variations, whose effect is generally stronger than that of demographic variations [25]. However, without demographic stochasticity the system never reaches the zero-abundance state, so in these theories one must introduce an arbitrary threshold below which a population is considered extinct. In section B of Supplementary Materials we show that our formula converges to the expressions obtained in the literature when demographic stochasticity is neglected, *if* the extinction threshold is chosen at

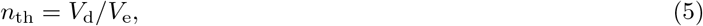

i.e., at the abundance level at which the strength of the demographic stochasticity becomes equal to that of the environmental stochasticity. This result clarifies the range of parameters for which infinite population models are applicable: the total population has to be much larger than *n*_th_. Furthermore, by introducing an absorbing boundary at this threshold value *n*_th_, one may obtain quantitative approximations (for quantities like the mean time to extinction) from models that do not take demographic stochasticity into account.

Our invasion formula has a few limitations. First, to derive it we assumed that the dwell time *δ* of the environmental variations is smaller than the time required for a population to invade. In the opposite case when *δ* is larger than the typical invasion time [which is the “quenched” scenario of [37]], the actual value of *δ* becomes insignificant, since the environment typically remains fixed throughout the invasion (see section C of Supplementary Materials). Our formula does not apply in this regime, but it is not required either, since in this case one may simply use the well-known formula for the probability of invasion in a fixed environment [29, 30] and then find the overall chance of invasion by taking an appropriately weighted expectation value over all possible environmental states.

Second, as mentioned earlier, Eq. (2) holds only if the parameters 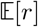, *V*_d_ and *V*_e_ are constant in the invasion regime. For this to happen the ratio *n*_f_/*N* must be small, otherwise density-dependent effects appear. Our method can still be used, but with the slight complication of having to integrate over the abundance *n* (see Eq. (15) in Methods). At the same time *n*_f_ cannot be so small that invasion occurs on timescales much shorter than the dwell time *δ*, otherwise the chance of invasion is determined by the initial conditions and the system is again in the “quenched” regime discussed above.

Third, to find the invasibility using abundance time series one must contend with sampling errors, which are common in ecological surveys [38]. As long as these errors are unbiased they do not affect 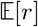, but they lead to an overestimation of *V*_e_. In some cases these errors may be filtered out using standard techniques [38, 39]. Otherwise, an informed guess about their effect on *V*_e_ is required. This problem actually highlights an important advantage of a closed-form formula like Eq. (2): knowledge of the functional dependence of Π on a quantity, such as *V*_e_, helps one to estimate the effect of errors in said quantity.

Species richness and genetic polymorphism reflect the balance between the rate of extinction and the rates (colonization, mutation, speciation, etc.) at which new types get established in the community. Invasibility, the chance of an invasion attempt to succeed, plays a crucial role in both kinds of processes. The rate of colonization is determined by multiplying a basal “attempt rate” (like the rate of mutation or speciation or migration from a regional pool) by Π. Similarly, given an attempt rate for extinction (the frequency with which a population visits a given low-density abundance), the number of independent attempts until extinction is distributed geometrically with mean 1/(1 − Π), so Π renormalizes the extinction rates. All in all, the parameter calculated here plays a key role in the main processes that govern biodiversity. Consequently, our formula should facilitate a better assessment of these important aspects of life science systems.

## Acknowledgements

This research was supported by the ISF-NRF Singapore joint research program (grant number 2669/17).

## Methods

### A. Derivation of Eq. (2)

In this paper we define the chance of invasion Π_*n*_ as the chance of a population of *n* individuals to reach a stated abundance *n*_f_ before extinction. Here we provide a general sketch of the derivation of this formula; a detailed derivation is presented in section A of Supplementary Materials.

Π_*n*_ satisfies the Backward Kolmogorov Equation,

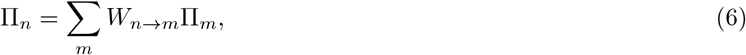

with the boundary conditions Π_0_ = 0 and 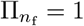. Here *W*_*n→m*_ is the transition probability: the chance to go from a state with *n* individuals to a state with *m* individuals during *δ* generations. Using the logarithmic parameter *z* = ln *n*, this equation translates to

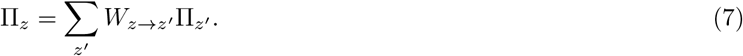

The conversion of *n* (and *m*) to *z* (and *z′*), and vice versa, is trivial, and in what follows we switch between the *n*- and *z*-notations as convenient.

As explained in the main text, our calculation requires the values of 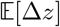, *V*_e_ and *V*_d_ (respectively the mean increase in *z*, the strength of environmental stochasticity, and the demographic stochasticity parameter) per dwell time *δ*. The extraction of these parameters from empirical or numerical time series is explained in Fig. 1 and the discussion around it. For the calculation of the invasion probability we assume that *n*_f_ « *N*, so that these three parameters are *n*-independent in the relevant regime. At the end of this section we refer to the case when this assumption does not hold. We also assume that the mean time to invasion is much larger than the correlation time of the environment; otherwise the timing of an invasion attempt determines its chance to succeed.

Given Eq. (1), the mean step size (∆*z*, during the dwell time *δ*) is 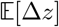, the second moment is 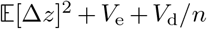 and the standard deviation of ∆*z* is 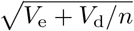. As discussed in section A of Supplementary Materials, our technique is based on the two-destination approximation, wherein we replace the actual random process (Eq. (7)) by a simpler process, a two-destination walk that preserves the first two moments of each jump.

Suppose that the appropriately-chosen jump size (either left or right) of the two-destination walk is 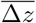, the probability of a right jump is *α* and that of a left jump is 1 − *α*. To keep the first two moments of this two-destination walk the same as the first two moments of the actual process, the parameters *α* and 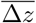 must satisfy

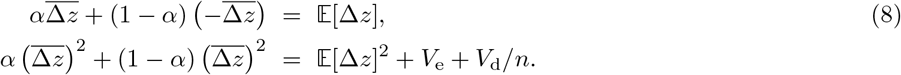

These equations give

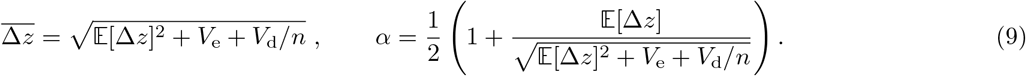

Accordingly, Eq. (7) becomes

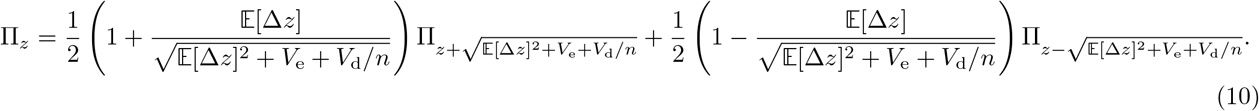

Applying our WKB-based approach, we express Π_*z*_ as 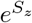, where *S*_*z*_ is assumed to be an adequately slowly varying function of *z* such that *S*_*z*+*dz*_, for a small change *dz*, may be approximated by 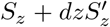, where a prime denotes a derivative with respect to *z*. For brevity of notation we use 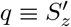. With this prescription, Eq. (S3) gives

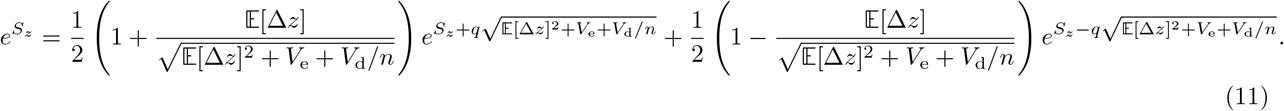

This yields a transcendental equation for *q*,

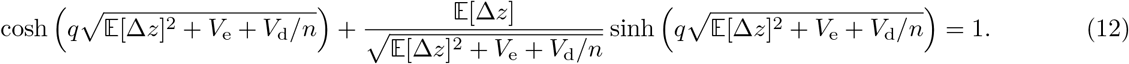

Remarkably, Eq. (S6) can be solved exactly,

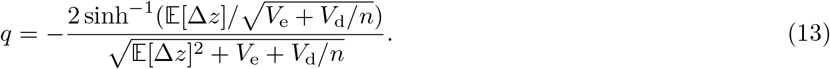

In section A of Supplementary Materials we provide a detailed exposition of all these steps. We also discuss there the relationship between the two-destination scheme with symmetric jumps (i.e., the same jump size 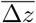 in the left and right directions), as utilized here, and the asymmetric-jumps scheme (with different jump sizes in the two directions) that we employed in [28].

Once *q* is obtained, an integration over it yields *S*_*z*_ = ∫*q*(*z*)*dz* = ∫[*q*(*n*)/*n*]*dn* (since *z* = ln *n*), and thence Π(*z*).

Eq. (S6), like Eq. (S1), admits also a trivial solution Π_*n*_ = constant, because the transition probabilities must sum to 1. As a result, the general solution for the chance of invasion is

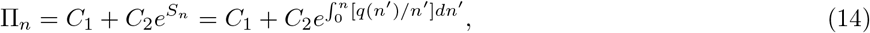

where the constants have to satisfy the boundary conditions Π_0_ = 0 and 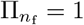. Hence we get

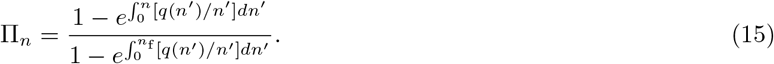

An analytical integration of *q/n*, where *q* is defined in Eq. (13), is not a simple task. To overcome this we employ the trick used in [28]. We note that *q* converges to 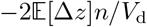 when *n* → 0 and to

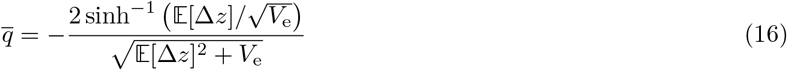

when demographic stochasticity is negligible (so *V*_d_/*n* is much smaller than 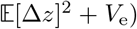. A good (and integrable) approximation for *q*(*n*) may be written down as

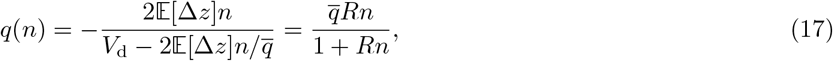

where *R* is defined in Eq. (4). This approximation becomes exact when *q* is small, while for larger values of *q* it interpolates between the correct expressions in the large-*n* and small-*n* regimes. Integrating *q*(*n*)/*n* over *n*, using the expression in Eq. (17), one obtains Eq. (2) of the main text from Eq. (15).

Extending this procedure to more general situations (e.g., when *n*_f_ is a finite fraction of *N*, so due to density-dependent effects 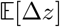, *V*_d_ or *V*_e_ vary across the invasion range 1 ≤ *n* ≤ *n*_f_) requires one to perform the integration in Eq. (15) over multiple *q*(*z*) regions, as described in [28]. This can be done either numerically or, in some cases, analytically.

### B. Details of the example models used

#### 1. Lottery model: discrete-time version

The two-species lottery model of Chesson and Warner [22, 36] is a simple and generic model of ecological dynamics. Its original, discrete-time version generalizes the classical Wright-Fisher model by allowing generations to overlap. Many other ecological models (e.g. Beverton-Holt) can be seen as modifications of the lottery model that account for more realistic processes (age structure, different dynamics for adults and seeds, etc.) but share the same basic structure.

In coexistence theory the lottery model has a particular importance as it exhibits the storage effect [stochasticity-induced stabilization; see a review in [21]]. In contrast with the naive viewpoint according to which increased environmental and abundance fluctuations decrease the coexistence time and cause more extinction events, stochasticity may facilitate coexistence under lottery dynamics.

The original formulation of the lottery model neglects demographic stochasticity. In models without demographic stochasticity, a population that starts from a positive abundance can never reach an abundance of strictly zero. This leads to an artificial necessity of having to introduce an arbitrary threshold to define extinction [22]. Here [as in [20]] we extend the lottery model to explicitly include demographic stochasticity, thus making the model more faithful to real-world situations.

The dynamics take place as follows. At the beginning of each timestep *t*, the community has *n*_*t*_ invading-species individuals and *N* − *n*_*t*_ resident species individuals (with *n*_*t*_ « *N* initially). Each individual produces a large number of seeds (or larvae, etc.), and the fitness of a species is related to the mean number of seeds produced by an individual. If an invader species individual produces exp(*s*) seeds per one seed of the resident species, the fitness ratio is exp(*s*) : 1. In a stochastic environment, *s* depends on *t*. To simulate it, we picked, at each step, a number *s*_*t*_ from a normal distribution with mean *s*_0_ and variance *σ*^2^.

After the seed-production step, any individual in the community dies with a probability *δ*, leaving on average *Nδ* empty gaps to be recruited. Seed dispersal is practically infinite, so spatial effects play no role (i.e., the dynamics are well-mixed). Therefore, the chance 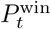 of the invader species to recruit any given gap is dictated by its share in the seed bank, namely,

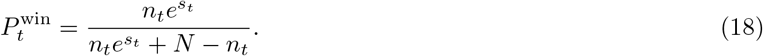

Accordingly, in each step the mean number of invaders satisfies 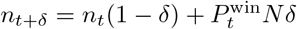.

Demographic stochasticity in this system has two sources. First, the number of invader species individuals that die in each step is not precisely *n*_*t*_*δ*, but is picked from a binomial distribution whose mean is *n*_*t*_*δ*. Analogously, the number of resident species deaths is distributed binomially around (*N* − *n*_*t*_)*δ*. Second, for a given number of gaps *G* (arising from the death of some invader and some resident species individuals) the number recruited by the invader species is again distributed binomially around 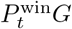.

#### 2. Lottery model: continuous-time (Moran) version

In the discrete-time version of the lottery model, as described above, the dwell time of the environment is simply *δ*, so there is a connection between the intrinsic dynamics of the population and the dynamics of the environment. To lift this restriction one may consider a continuous-time version of the lottery model [21, 40].

In this version, in each elementary step, only one individual is chosen at random to die. The resulting gap is recruited by an offspring of the focal species with probability 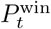 [Eq. (18)] and by an offspring of the resident species with probability 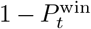. In each elementary step, the environment switches with probability 1/(*Nδ*), so the persistence time is picked from a geometric distribution whose mean is the dwell time *δ*.

An analytic derivation of the parameters 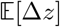, *V*_e_ and *V*_d_ for this Moran model is presented in section E of Supplementary Materials.

#### 3. Forest dynamics model

The forest dynamics model, as employed by [18, 19], is based on Leslie-Gower dynamics. Growth is divided into a sapling stage and an adult stage. The sapling dynamics are described by

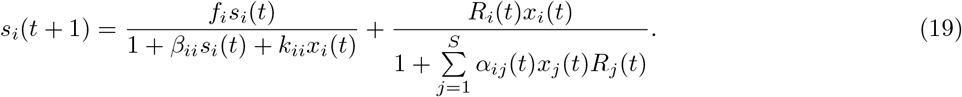

Here the subscripts (*i* or *j*) label the species and *t* denotes the time, measured in years. The parameter *s* stands for the sapling density, *f* represents the sapling fraction surviving from a year to the following one, *R* denotes the rate at which seeds (or seedlings) are generated, and *x* denotes the frequency of the adults. *S* is the total number of species in the system, and *β*, *k* and *α* quantify the competition faced by the saplings (of a given species) from other saplings of their own species and from the adults and the saplings of the other species.

The two terms on the right hand side in Eq. (S22) correspond to the saplings surviving from previous years, and to new saplings that are generated by the adults. In order to focus on the effect of environmental variability, we have taken the values of the competition parameters to be *α_ij_* = 1 and *β_ii_* = *k_ii_* = 0 for all *i, j* in our simulations, mimicking the approach employed by [18, 19].

The adults in the model follow dynamics resembling the lottery model [36],

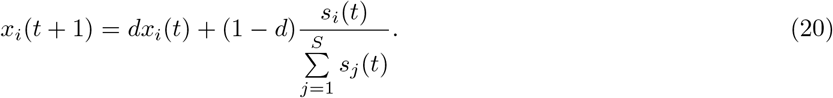

Here *d* denotes the fraction of adults of a given species that survives in each timestep. The ratio of the frequency of the saplings of a species to the total number of saplings across all species determines the probability of that species to take up an open adult gap. In the limit *f*_*i*_ = 0 for all *i*, the model loses its “memory” of the saplings surviving from previous years and converges to the lottery model. It is due to this long-term memory that in the forest dynamics model the dwell time *δ* of the environment is not directly related to the chance of an individual to die during *δ*, as in the two versions of the lottery model.

### C. Numerical simulations

We ran Monte Carlo simulations of the three models described above in order to check the predictions of Eq. (2).

In the discrete-time lottery model, the number of dying individuals from the invading species, which had an abundance *n*_*t*_ at the beginning of a timestep, was picked randomly from a binomial distribution with *n*_*t*_ trials and the chance of success *δ*. Similarly, the number of individuals dying from the resident species was picked from a binomial distribution with *N* − *n*_*t*_ trials and the chance of success *δ*. This meant that the average total number of deaths (across both the species) in each timestep was *Nδ*, as desired. If *d*_1,*t*_ and *d*_2,*t*_ denote the number of deaths thus suffered by the invading and the resident species, then the number of births for the invading species was picked randomly from a binomial distribution with *d*_1,*t*_ + *d*_2,*t*_ trials and the chance of success 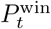. The resident species was allowed to fill up all the remaining empty gaps, thus preserving the total population as *N* individuals after every timestep.

We modified the procedure slightly to simulate the dynamics in the limit of infinite *N* (with *n*_*t*_ still finite). In this case we picked, as above, the number of dying invading species individuals from a binomial distribution with *n*_*t*_ trials and the chance of success *δ*. The number of births, however, of the invading species was picked from a Poisson distribution with mean 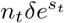 (with *s*_*t*_ again picked randomly from the values *s*_0_ ± *σ*). In this case the number of deaths and births of the resident species is immaterial.

The numerical results in Fig. 2 were not obtained from a Monte Carlo simulation but from an exact numerical solution for the chance of invasion, as described in section 4 of [28].

For the Leslie-Gower model, a detailed exposition of our simulation technique (including Matlab codes) and of the parameters used in these simulations is provided in section F of Supplementary Materials.

## Supplementary Material

### A. The two-destination approximation and the derivation of 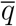

In the Methods section we discussed the derivation of our formula, Eq. (2). Here we would like to discuss it further, and in particular to explain the differences between the two-destination approximation used in [28] and the one used here.

The analysis of a stochastic system is based on transition probabilities. These transition probabilities are usually defined per elementary step, where an elementary step could be a single birth-death event, for example. When a system is subject to environmental fluctuations, the basic rates, and hence the transition probabilities, vary through time and change over timescales that are longer than a single elementary step. One cannot average the single birth-death transition probabilities over all environmental states, as this approach neglects temporal correlations (which result from adjacent steps tending to be in the same environment) and replaces the fluctuating environment by its average. Averaging over environmental states is allowed only when the elementary objects of the stochastic process are the transitions during the dwell time of the environment. In the corresponding Backward Kolmogorov equation,

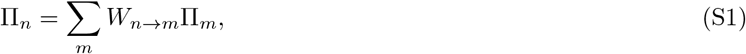

the *W*_*n→m*_ reflect transition probabilities during the dwell time *δ* and typically involve many destination states *m* for each *n*. The emerging problem is quite complicated to analyze.

The “jack of all trades” approach to this problem is the diffusion approximation, which is based on an expansion of the quantity of interest (Π, say) to second order in *n* − *m*. This procedure involves two levels of approximation. The first level is the continuum approximation, which depends on Π_*n*_ being a smooth function over the integers. Second, the second-order expansion implies that only the first two moments of *W*_*n→m*_ are important, so the mean and the variance (or the mean and the second moment) of the jump at each *n* determine the outcome of the process.

The failure of the diffusion approximation in some circumstances has been analyzed by many authors [32, 35, 41]. In [28] we pointed out that the main problem is with the first assumption, not the second: relying only on the first two moments yields accurate results as long as one avoids the continuum approximation. Therefore, we suggested an approach based on a large-deviations (controlling-factor WKB) technique that resolved the continuum approximation problem by assuming that the logarithm of Π, instead of Π itself, is smooth. We retained the second assumption, namely that the outcome depends only on the first two moments. This meant that the actual process could be replaced by an effective process in which instead of all *m* possible destination states there are only two destinations, properly chosen such that the mean *μ* and the second moment *ρ*^2^ of the jump, defined by

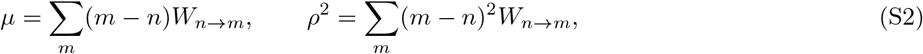

are preserved. Both *μ* and *ρ* are *n*-dependent, but for the sake of simplicity we omit the *n* index in this discussion.

There are infinitely many possible choices for this two-destination scheme. In general, for each *n* one may replace the actual full process by a process with only two jumps, from *n* to *m*_1_ with probability *α* and from *n* to *m*_2_ with probability 1 − *α*, where the only restriction on *m*_1_, *m*_2_ and *α* is that the resulting mean and second moment of the new process are the same as those of the original process. The scheme we adopted in [28] was based on equal jump probabilities (*α* = 1/2) and unequal jump lengths (|*m*_1_ − *n*| second moments equal *μ* and *ρ*^2^, respectively, this leads to:

| Destination | Probability | Jump length |
| --- | --- | --- |
| $m_1$ | $\alpha = 1/2$ | $\Delta m_1 = m_1 - n = \mu + \sqrt{\rho^2 - \mu^2}$ |
| $m_2$ | $1 - \alpha = 1/2$ | $\Delta m_2 = m_2 - n = \mu - \sqrt{\rho^2 - \mu^2}$. |

Clearly, the first moment of this walk is (∆*m*_1_ + ∆*m*_2_)/2 = *μ* and the second moment is [(∆*m*_1_)^2^ + (∆*m*_2_)^2^]/2 = *ρ*^2^.

Here we adopt an alternative scheme, in which the jump lengths are kept equal in magnitude and the jump probabilities are unequal, which leads to the following choice:

| Destination | Probability | Jump length |
| --- | --- | --- |
| $m_1$ | $\alpha = \frac{1}{2} + \frac{\mu}{2\rho}$ | $\Delta m_1 = m_1 - n = \rho$ |
| $m_2$ | $1 - \alpha = \frac{1}{2} - \frac{\mu}{2\rho}$ | $\Delta m_2 = m_2 - n = -\rho$ |

It may be verified that again, *α*∆*m*_1_ + (1 − *α*)∆*m*_2_ = *μ* and [*α*(∆*m*_1_)^2^ + (1 − *α*)(∆*m*_2_)^2^] = *ρ*^2^.

When only the two destinations *m*_1_ and *m*_2_, with their specified probabilities, are allowed, under the old scheme of [28] (*α* = 1/2), Eq. (S1) is reduced to

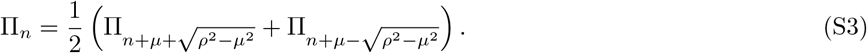

With the new two-destination scheme (equal jumps) used here, Eq. (S1) leads likewise to the BKE equation

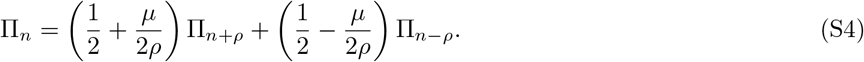

Under the WKB approach, we make the ansatz

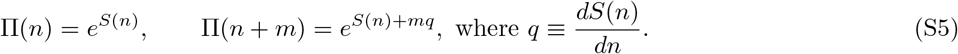

When plugged into Eq. (S3), this ansatz results in the transcendental equation

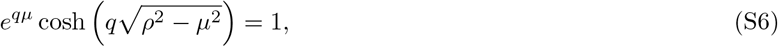

and when plugged into Eq. (S4), the WKB ansatz leads to

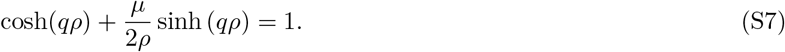

Since Eq. (S6) (which was employed in [28]) cannot be solved analytically (to our knowledge), in [28] we used the approximate solutions

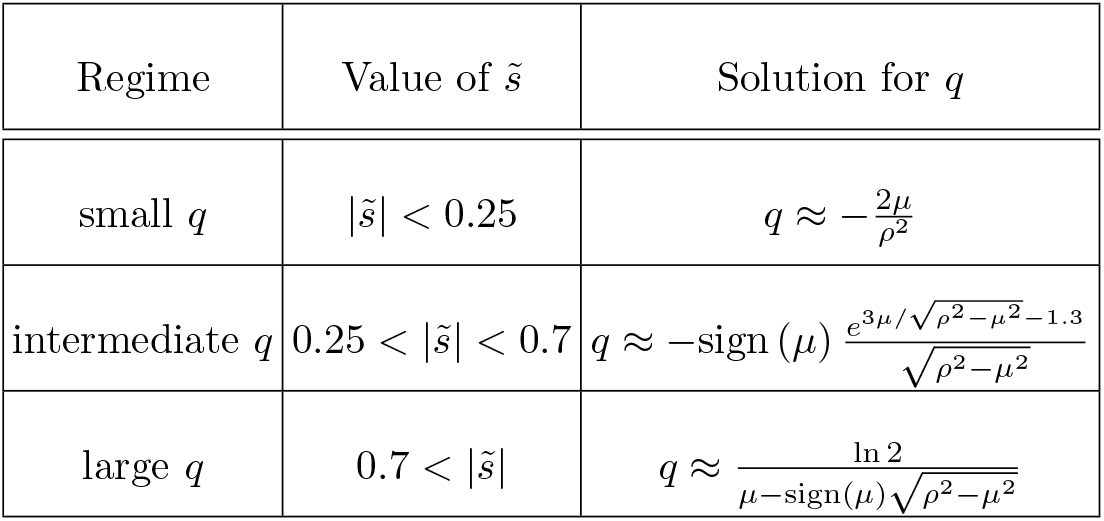

where 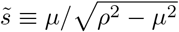.

In contrast, Eq. (S7) does have an analytical solution,

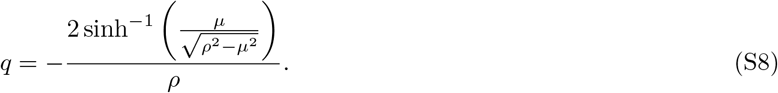

This is the solution that we have used throughout the main text of our present paper, only with the notational change of the mean jump length *μ* being replaced by 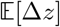 and *ρ* being replaced by 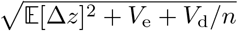. In the main text we have also mostly worked in *z*-space, where *z* = ln *n* is the logarithm of the abundance *n*. It is easy to switch from *z*-notation to *n*-notation and vice versa, as convenient.

What are the differences between the two schemes of choosing the two destinations? Is there a “correct” scheme?

First, we stress that the two expressions for *q*, in Eq. (S8) and in the solutions presented in the table above, coincide in the small-*q* regime, where stochasticity is strong. They differ mainly in the large-*q* regime when 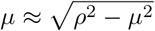, i.e., when the mean value of the jump length is approximately equal to its standard deviation 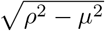. In this regime the scheme we have used in [28] yields a diverging value for *q*, while the expression in Eq. (S8) is finite.

The origin of this difference is easy to understand. When one uses the old “equal probabilities-unequal jump lengths” scheme, the length of the shorter jump is the mean minus the standard deviation. When the mean is larger than the standard deviation both jumps are in the same direction, so the abundance either grows in every step or diminishes in every step. Thus the chance of fixation is either strictly 0 or strictly 1, which manifests itself in the divergence of *q*. In contrast, under the new “equal jump lengths-unequal jump probabilities” scheme, jumps are always in opposite directions and the bias *μ* affects only their chances (the *α*-values), so for any strength of the bias there is still a chance (even if small) for both extinction and fixation.

Which of the two schemes is more suitable depends on the statistics of the fitness variations. If the possible deviations of fitness values from their mean are limited (in other words, if the probability distribution function from which fitness fluctuations are drawn is compact), then once *μ* gets large enough the actual process will move deterministically towards either fixation or extinction. The correct scheme, then, is the one used in [28], i.e., with equal jump probabilities and unequal magnitudes of the jump length, which yields the transcendental equation (S6) in which *q* diverges when ∆*m*_1_ and ∆*m*_2_ have the same sign. On the other hand, when fitness fluctuations are unlimited, even with low probability (when the probability distribution function is non-compact, e.g., Gaussian) then the scheme used in the current paper, with equal magnitudes of the jump length and unequal jump probabilities, is better. This is so because in this case there is always a chance for movement in both directions, towards extinction and fixation. The appropriate expression for *q* then is the one in Eq. (S8), the solution of Eq. (S7).

### B. Limits of Eq. (2) of the main text

In this section we consider the limits of the chance of invasion (Eq. (2) of the main text) when the environment is fixed and when the effect of demographic stochasticity is taken into account only as a threshold density below which a population is considered extinct. The results for a fixed environment are known, and we show that our general expression converges to these results. Theories without demographic stochasticity usually introduce an arbitrary extinction threshold, and our expression allows one to define the appropriate threshold value.

#### B1. Fixed environment

In a fixed environment (*V*_e_ = 0), invasion reflects the interplay between 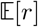 and demographic stochasticity. In this case the term *V*_d_/*n* in Eq. (13) of the main text must be kept for all values of *n* (unlike when *V*_e_ > 0, in which case *V*_d_/*n* can be ignored for large enough *n*; see next subsection), so calculations of the chance of invasion must rely on Eq. (15) of the main text.

Plugging Eq. (13) (in the limit where *V*_e_ = 0) into the integral in the numerator of Eq. (15), one gets

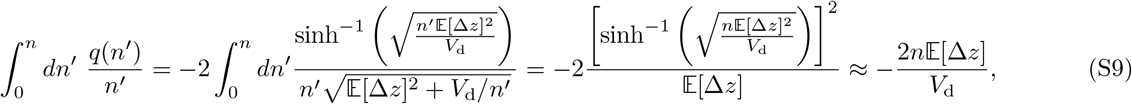

where the last approximation holds when the argument of the sinh^−1^ function is small. The chance to reach *n*_f_ before extinction is, thus,

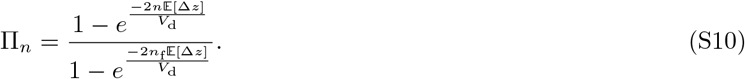

The demographic parameter *V*_d_ is 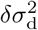, where 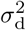 is the variance of the number of offspring that each individual produces during its lifetime [25]. Thus *V*_d_ = 2*δ* for a neutral 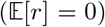 continuous-time (Moran) process, for which the lifetime probability for *m* offspring is 1/2^*m*+1^, and *V*_d_ = *δ* for a neutral Wright-Fisher process, where *m* is picked from a Poisson distribution with mean 1. *V*_d_ is very close to these values even if the selective forces are not strictly zero but are weak 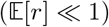.

When *n*_f_ » 1 and 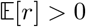, the denominator of Eq. (S10) is very nearly 1, therefore the chance of a single mutant (*n* = 1) to reach high abundances depends only on the numerator. Accordingly, in a Moran process the chance of establishment for a single slightly beneficial mutant is approximately 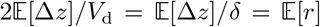 while under Wright-Fisher dynamics it is 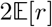, in agreement with known results that were obtained long ago using branching process analysis [29] or the diffusion approximation [30]. Our technique makes it possible to obtain higher-order corrections to these expressions.

#### B2. Replacing demographic stochasticity with absorbing boundary at a given abundance threshold

When a population is large, the effect of demographic stochasticity (which reflects the differences in reproductive success and dying rates of different individuals) is much weaker than the effect of stochastic environmental variations [25]. Mathematically, the amplitude of demographic fluctuations scales with the square root of the population size, while fluctuations associated with environmental stochasticity scale linearly with the population size. Therefore, many theories of community dynamics, permanence, and stability properties do not take demographic stochasticity into account [5–8].

However, demographic stochasticity reflects the quantization of the number of individuals in a population. Without demographic stochasticity even an exponentially decreasing population never reaches zero. As a result, while in reality a population can go extinct, in theories without demographic stochasticity strict extinction is impossible and, as its environment varies, a population may always recover from long periods of decline. To account for this problem, theories without demographic stochasticity define a threshold value below which a population is considered extinct [8, 22]. The choice of this threshold is arbitrary. This does not pose a problem for theories of permanence and uniform stability, because these theories are focused on a binary distinction between stable and unstable systems, which does not depend on the exact location of the threshold. However it may be very instructive to make these theories quantitative or semi-quantitative by an informed choice of this threshold. In this subsection we compare our results with the outcome of a theory without environmental stochasticity, and determine such an appropriate choice.

When demographic stochasticity is neglected, the invader population follows the dynamics of an asymmetric random walk along the *z*-axis. If, additionally, the diffusion approximation holds, then the chance of invasion satisfies the biased diffusion equation,

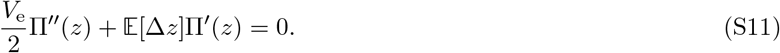

The solution of Eq. (S11), with the boundary conditions Π(*z*_th_) = 0 and Π(*z*_f_) = 1, gives the chance of a random walker at *z* to reach a given destination *z*_f_ > *z* before reaching some lower threshold value *z*_th_ < *z*. This solution takes the form [42]

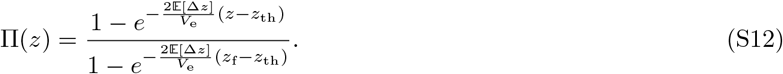

This well-known result was applied to population dynamics in [21] [their Eq. (46)].

How can one obtain this expression from Eq. (2) of the main text? Upon taking the limit *V*_e_ » *V*_d_ and 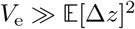, one finds

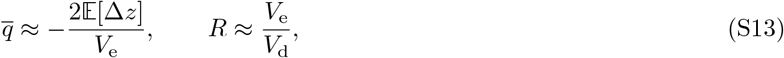

so Eq. (2) takes the form

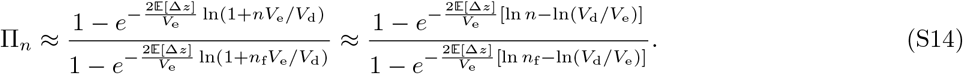

Writing *z* = ln *n* and *z*_f_ = ln *n*_f_, one realizes that the two expressions Eq. (S12) and Eq. (S14) coincide if

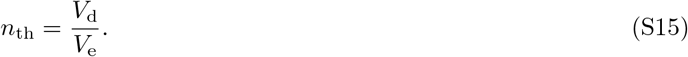

The correct threshold, thus, appears at the abundance level at which the strength of the demographic stochasticity becomes equal to that of the environmental stochasticity. Demographic stochasticity, which is unbiased along the arithmetic-abundance axis, generates a bias towards extinction on the log-abundance axis [25, 43]. At the threshold point, when its effect is dominant, the population sticks to the low-abundance extinction regime.

### C. Supplements for Fig. 2 of the main text

In Fig. 2 of the main text we presented the chance of invasion Π_*n*=1_, as obtained from numerical solutions of the discrete-time lottery model, for different values of the dwell time *δ* and the amplitude of fitness variations *σ*. We plotted Π_*n*=1_ against the invasion parameter 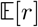 and obtained scattered results, without a specific trend associated with the mean growth rate. When plotted against the predictions of Eq. (2), we obtained a nearly perfect data collapse. In Fig. 2 we used *n*_f_ = 200. Here we present corresponding figures for larger and smaller values of *n*_f_.

For *n*_f_ = 500, the data-collapse is even better (if only a little) than the results obtained for *n*_f_ = 200, as seen in Figure S1. When *n*_f_ = 20 (Figure S2) some deviations appear.

**FIG. S1:**
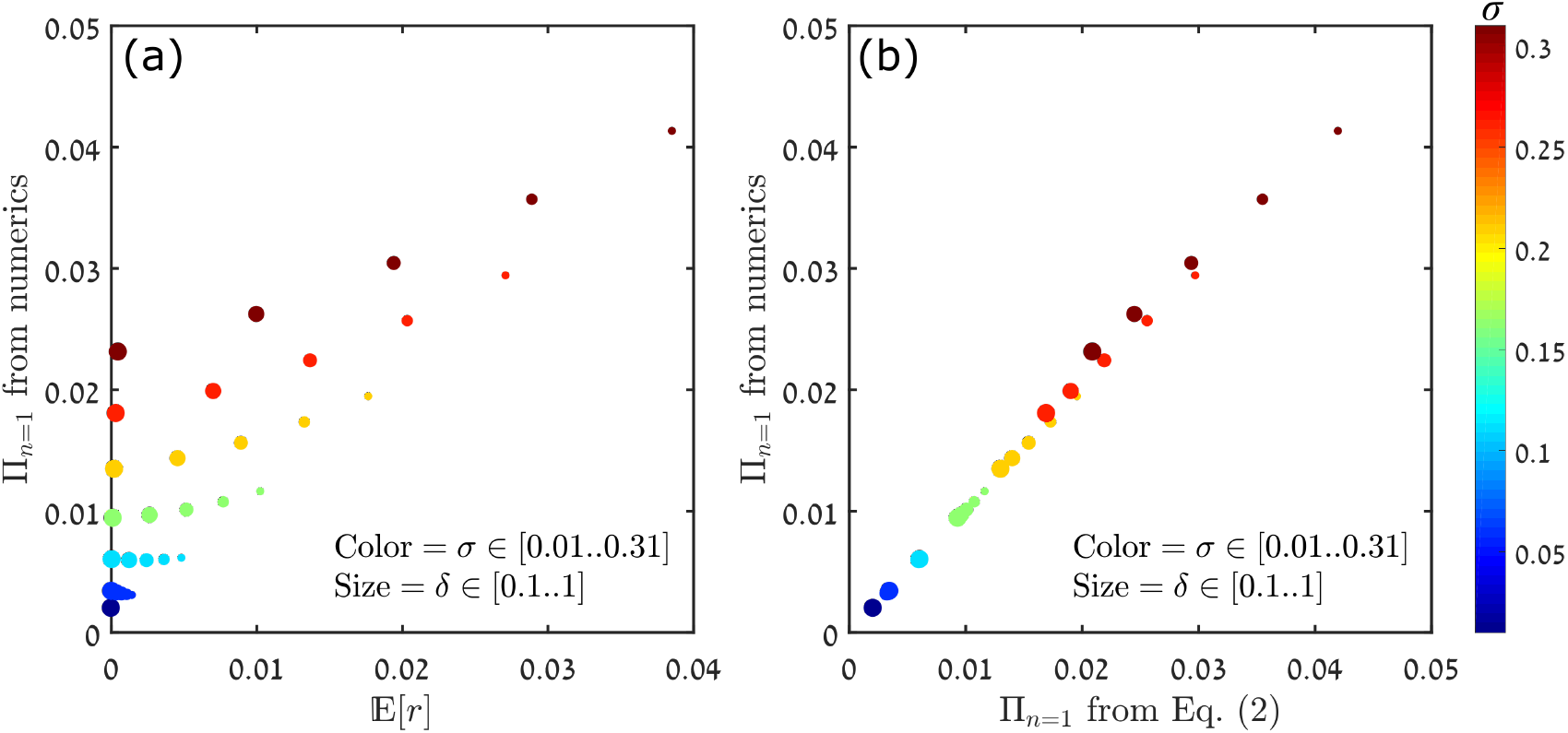
The chance of invasion for *n*_f_ = 500. All other parameters, as well as the size code for *δ*, are identical to those used in Fig. 2 of the main text.

Figs. S1 and S2 indicate that as *n*_f_ decreases, the parameter *δ* diminishes in importance. This is so because in the extreme case, as *n*_f_ becomes very small, there is no dependence of the chance of invasion on *δ*: a population reaches the invasion point, or goes extinct, before the environment flips. As explained in the main text, this corresponds to the “quenched” regime of Mustonen and Lässig [37], which is not covered by our theory. Even so, Fig. S2 suggests that the disagreement is not large even for *n*_f_ as small as 20.

The results presented in Fig. 2 of the main text, and correspondingly in Figs. S2 and S1 here, were obtained for a model with *N* → ∞. This allows us to solve analytically for the transition probabilities and hence to obtain the chance of invasion from an exact numerical solution of the linear problem posed by the Backward Kolmogorov equation (S1). The results presented in Figures 3 and 4 of the main text were obtained for finite-*N* systems via Monte-Carlo simulations of the elementary process.

**FIG. S2:**
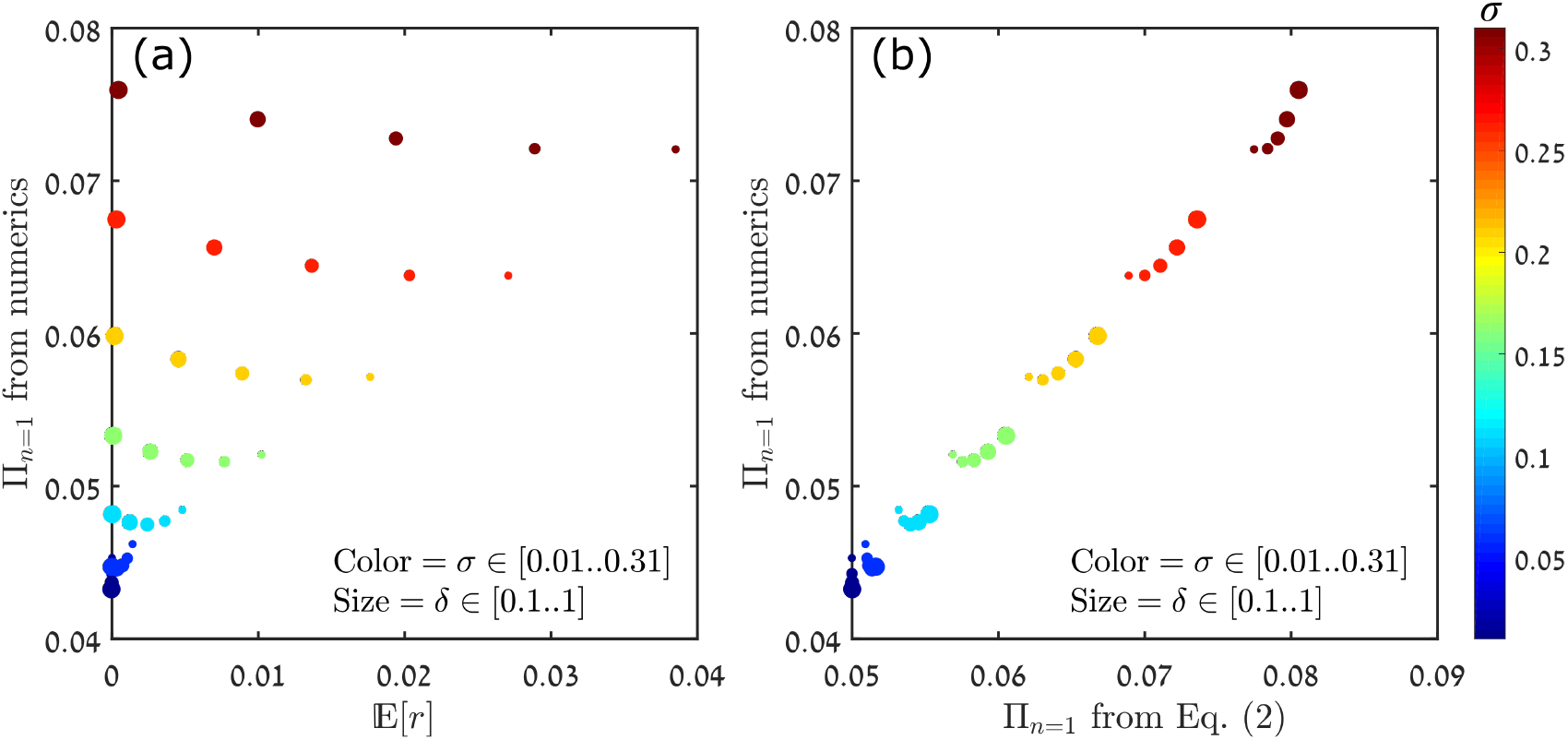
The chance of invasion for *n*_f_ = 20. All other parameters, as well as the size code for *δ*, are identical to those used in Fig. 2 of the main text. As *n*_f_ decreases the dependence of Π on *δ* reduces and becomes non-monotonic.

### D. Supplements for Fig. 3 of the main text

In panel (a) of Fig. 3 of the main text we showed the chance of invasion in the Moran model as a function of the starting population *n* for different values of the fitness advantage *s*_0_, where *σ* = 0.3, *δ* = 0.1, *N* = 5000 and *n*_f_ = 200. In this figure, as in Fig. 4 of the main text, the results were obtained for finite-*N* systems via Monte-Carlo simulations of the elementary process. Here we show the same figures for a few other values of *N* and *n*_f_.

The differences between the plots for *N* = 5000 (Fig. 3(a) in the main text) and *N* = 10000 (Fig. S3 here) are minute. The agreement of the theory with the simulations is slightly better when *N* = 10000, because the ratio *n*_f_*/N* is smaller, so density-dependent effects in the invasion regime are even weaker.

**FIG. S3:**
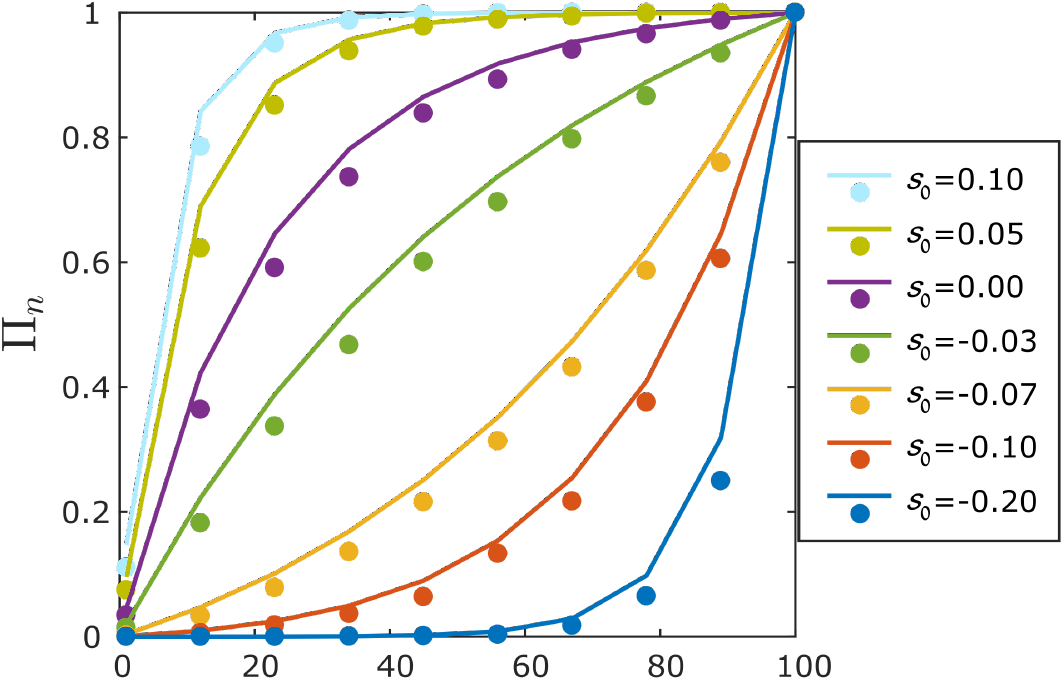
The chance of invasion as a function of the initial abundance *n*, with *N* = 10000. All other parameters are identical to those used in Fig. 3(a) of the main text.

In the same vein, the agreement between the simulation results and the theoretical formula in Eq. (2) of the main text improves further (even if minutely) when *N* is increased to a very large value with *n*_f_ held fixed (so that density-dependent effects are negligible). Fig. S4 shows an example of such a case, with *N* = 10^7^ and *n*_f_ = 100.

On the other end, as *N* decreases (and *n*_f_ is held constant), density-dependent effects become noticeable and the agreement between our formula and the numerical results starts to get poorer. For example, for *N* = 1000 (Fig. S5), the deviations are a little more pronounced than those observed in Fig. 3 of the main text.

However, if *n*_f_ and *N* are decreased proportionately, density-dependent effects decline again and the agreement between theory and numerics improves, as seen for *N* = 1000*, n*_f_ = 50 (Fig. S6).

**FIG. S4:**
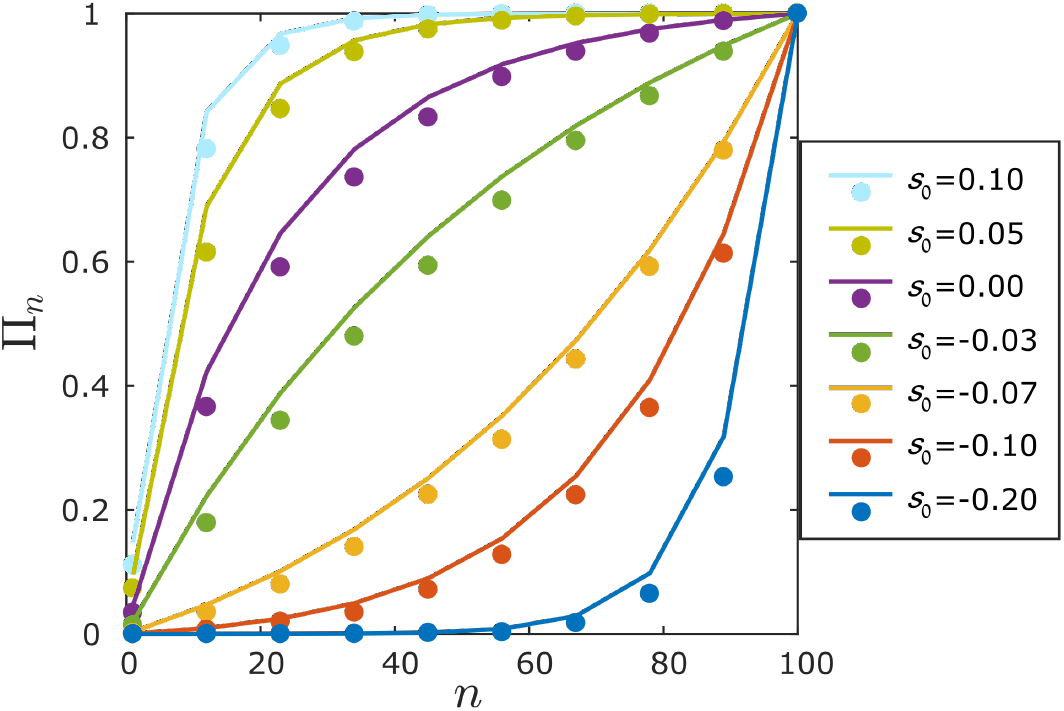
The chance of invasion as a function of the initial abundance *n*, with *N* = 10^7^ and *n*_f_ = 100. All other parameters are identical to those used in Fig. 3 of the main text.

**FIG. S5:**
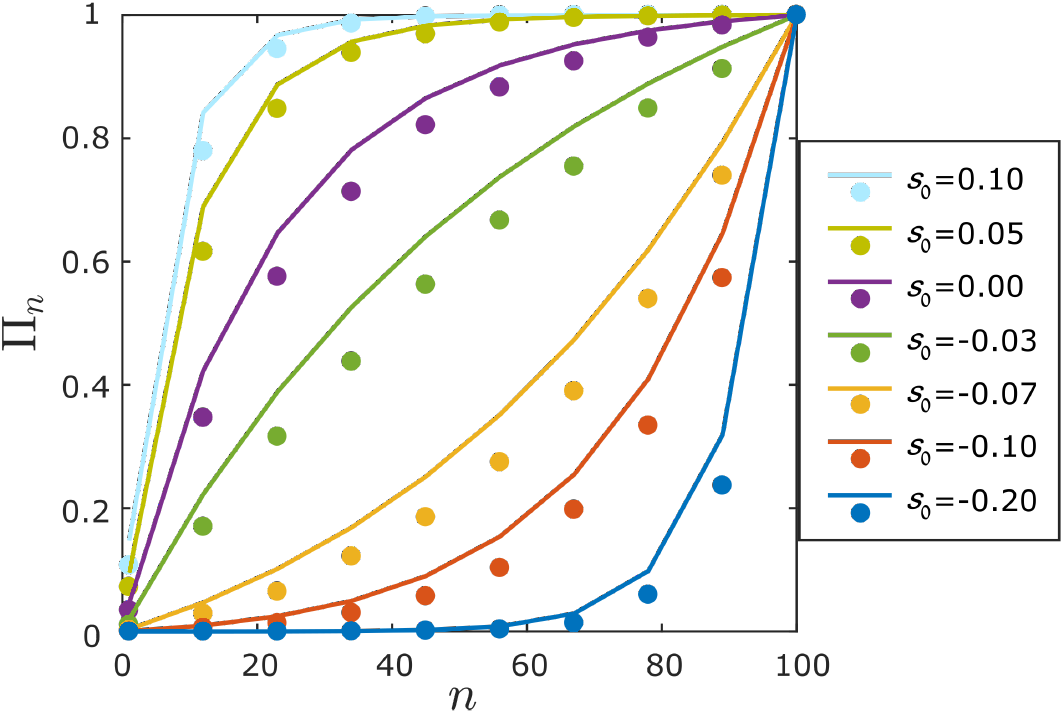
The chance of invasion as a function of the initial abundance *n*, with *N* = 1000 and *n*_f_ = 100. All other parameters are identical to those used in Fig. 3 of the main text.

**FIG. S6:**
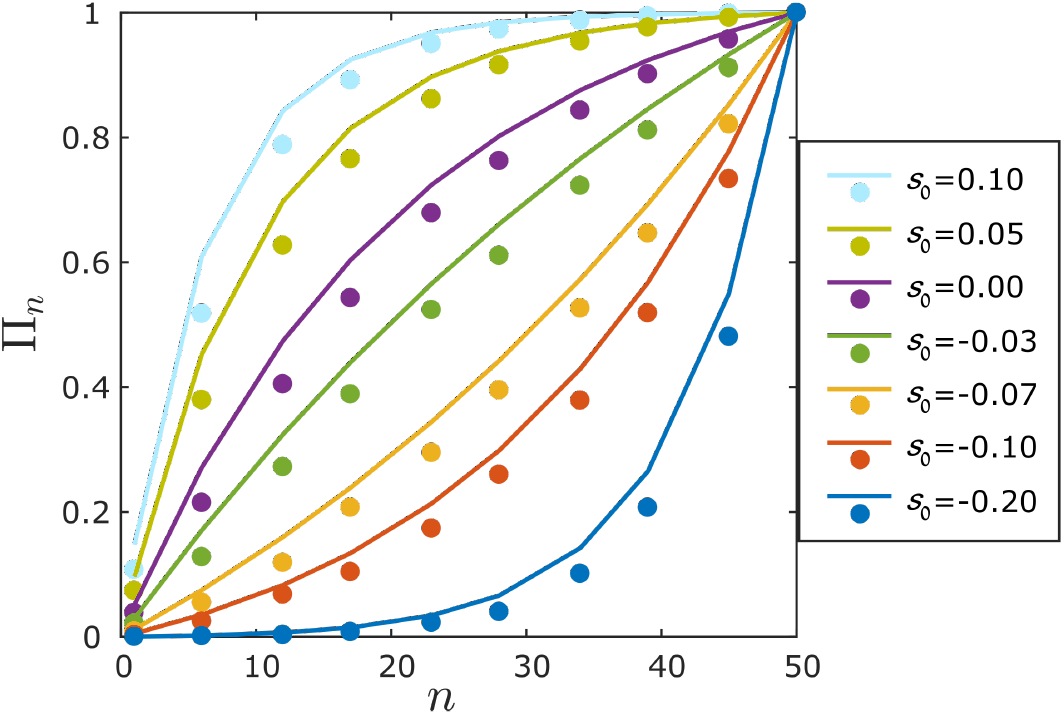
The chance of invasion as a function of the initial abundance *n*, with *N* = 1000 and *n*_f_ = 50. All other parameters are identical to those used in Fig. 3 of the main text.

### E. Invasion parameters in a lottery model with continuous-time (Moran) dynamics

Our formula, Eq. (2) of the main text, employs three quantities, the mean value 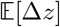 of the growth rate, its variance *V*_e_ and the demographic stochasticity parameter *V*_d_, with all these parameters calculated over the dwell time *δ*. In the main text (Figure 1) we presented a general method aimed at calculating these quantities from empirical or from numerically-generated time series. Here we would like to present a case in which these parameters may be calculated analytically given the model details. This sheds some light on the relationship between the elementary parameters of a system and the three parameters used in Eq. (2), and clarifies, in particular, how these quantities were obtained for Fig. 3.

The Moran model is presented in the Methods section. In each elementary step a single individual is picked at random to die, and its place is recruited by an individual of the focal species with probability *P*^win^ (see Eq. (18) of Methods), otherwise it is recruited by the rival species. *P*^win^ depends on the environmental conditions (say weather), which are held to change, on average, every *δN* elementary birth-death events (i.e., every *δ* generations, with the time of an elementary event being 1/*N* generations). The weather conditions are manifested in the exponential factor which appears in the fitness of the focal species, 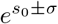, where *s*_0_ is the fitness advantage of the focal species and ±*σ* is the effect of the environment. (To simplify the calculations we assume here a dichotomous noise. The interested reader may read up the relationship between this case and that of more general forms of the noise in [44].)

Let us initially calculate the change in abundance without including the demographic stochasticity. We will later add it in the appropriate place. With the setup described above, the change in the frequency of the focal species *x* = *n/N* in each elementary event is:

| Case | Probability |
| --- | --- |
| $x \rightarrow x + \frac{1}{N}$ | $\frac{x(1-x)e^s}{1-x+xe^s}$ |
| $x \rightarrow x - \frac{1}{N}$ | $\frac{x(1-x)}{1-x+xe^s}$ |
| $x \rightarrow x$ | $1 - \frac{x(1-x)(e^s+1)}{1-x+xe^s}$ |

Here *s* = *s*_0_ ± *σ* is the instantaneous selection (log-fitness) parameter. The chance for the dying individual to be picked from the non-focal species is 1 − *x*, and the chance for the focal species to then take over its slot is *xe^s^*/(1 − *x* + *xe^s^*), giving the above probabilities for *x* to increase by 1/*N* and to decrease by 1/*N*.

For invasion we have *x* « 1 and *xe^s^* « 1 (the latter inequality being true, for any *s*, if *N* is large enough). Upon replacing 1 − *x* by 1 in the probabilites to increase and decrease by 1/*N*, and then subtracting the sum of these from 1 in order to find the probability to retain the same abundance, we find that the probabilities simplify to:

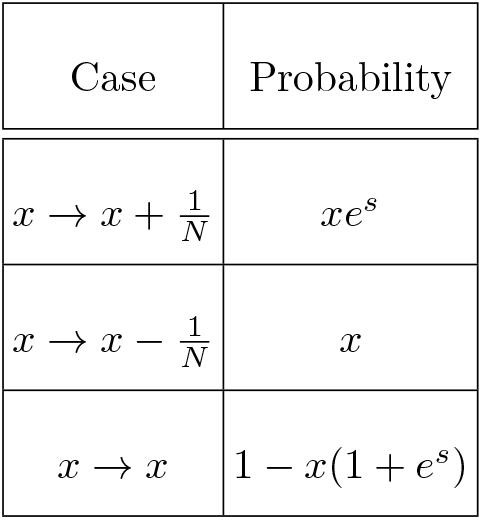

Therefore, the mean change in *x* in one elementary event, (*dx/dt*)/*N*, is

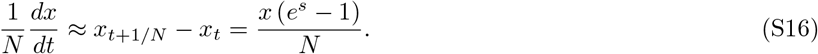

Integrating this quantity over the dwell time *δ* one obtains

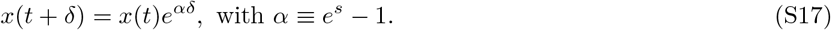

The demographic stochasticity in this system is 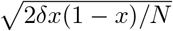, meaning that the net value of *x*, including demographic fluctuations, is (again with *x* « 1, and with demographic stochasticity represented by dichotomous demo-graphic noise)

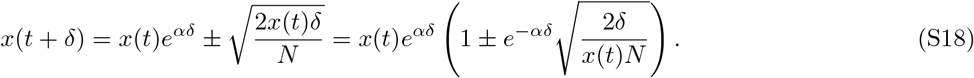

Defining *z* = log *n* = log *xN*, we have

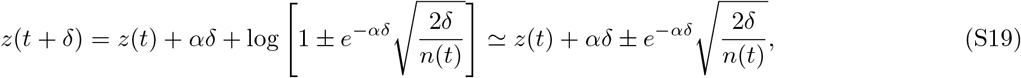

where *n*(*t*) = *x*(*t*)*N*.

To be able to use our formula in Eq. (2) in the main text we need the mean value 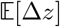 of ∆*z* and its variance Var[∆z]. From Eq. (S19), putting in the two possible values of *α* as 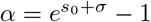 and 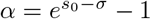, we find

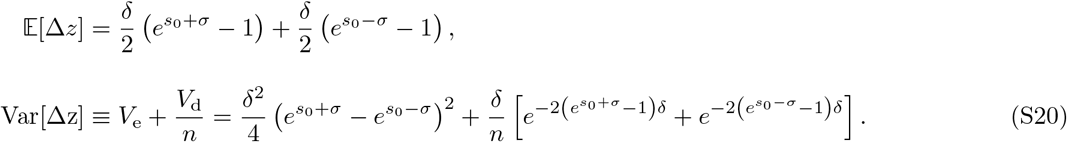

Therefore, we get our desired quantities,

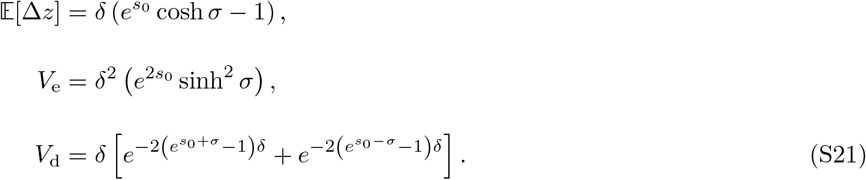

Putting these quantities into Eq. (2) of the main text, we obtained the analytical prediction for the chance of invasion in the Moran model, which was used in Fig. 3.

It is important to distinguish *σ* from *V*_e_: the first is the amplitude of *fitness* (more accurately, log-fitness) variations whereas the second is the amplitude of environmentally-induced *abundance* fluctuations. The value of *σ* affects not only *V*_e_ but also 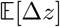 (and therefore 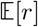) and *V*_d_. (The increase of 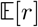 with *σ* is a manifestation of Chesson’s storage effect [36].) By and large, the relationship between the elementary parameters of the dynamics, such as *σ*, and the parameters of Eq. (2) may be complicated.

### F. Supplements for Fig. 4 of the main text

To demonstrate the effectiveness of our invasion formula, and the procedure involved in using it, we considered in Figure 4 of the main text a generic model of forest dynamics, namely the Leslie-Gower model, as employed in [18, 19], and the accompanying empirical time series for the recruitment rates of the two species *Spondias mombin* and *Spondias radlkoferi* as presented in [18]. Under the original parameters of [18] an invasion of one of the species is nearly certain, and occurs very fast, even for very small values of *n*. Therefore, in our demonstration we modified some of the parameters (namely, the survival probability of adults and of saplings, and the overall strength of recruitment fluctuations) while preserving the proportions and the cross-correlations of the original recruitment time series. Here we explain the procedure in detail and provide an annotated Matlab code for each step.

Our starting points are equations (20) and (21) of the main text. When the model is limited to two species (indexed by *i*) and the simplifications made in [18], as detailed in the Methods section, are adopted, the deterministic map for sapling dynamics is given by

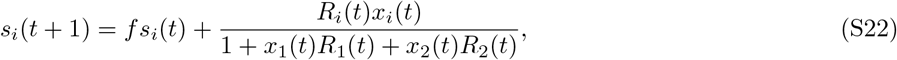

where *s*_*i*_(*t*) is the density of saplings of the species *i* at time *t*, *f* is the fraction of surviving saplings (of each species) from one year to the next, and *R*_*i*_(*t*) is the recruitment rate of species *i* at time *t*. Similarly, the deterministic map for adult dynamics is

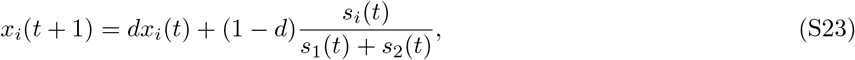

where *x*_*i*_(*t*) is the density of the adults of species *i* at time *t*, and *d* is the fraction of surviving adults (of each species) from one year to the next.

To obtain Fig. 4 of the main text, we first performed Monte-Carlo simulations of the above model, calculating from them the chance of a population of *n* individuals to grow in abundance to *n*_f_ before going extinct. In our simulations only the density of adults was quantized, i.e., demographic stochasticity entered the model only at the level of the adult dynamics. The sapling density was taken as a continuous variable, since the number of saplings is typically much larger than the number of adult trees. In subsection F1 we provide the Matlab code for this MC simulation. A typical outcome is presented in Fig. S7.

In order to use Eq. (2) of the main text one must find *δ*, 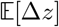, *V*_e_ and *V*_d_. To do that we generated a long time series of the same dynamics as described in Eqs. (S22) and (S23), using the code provided in subsection F2 below. We stipulated reflecting boundary conditions, so that *n* = 0 became *n* = 1 and *n* = *N* became *n* = *N* − 1. This was done to make it easier to obtain adequately long time series. Typical time series obtained are shown in Fig. S8.

**FIG. S7:**
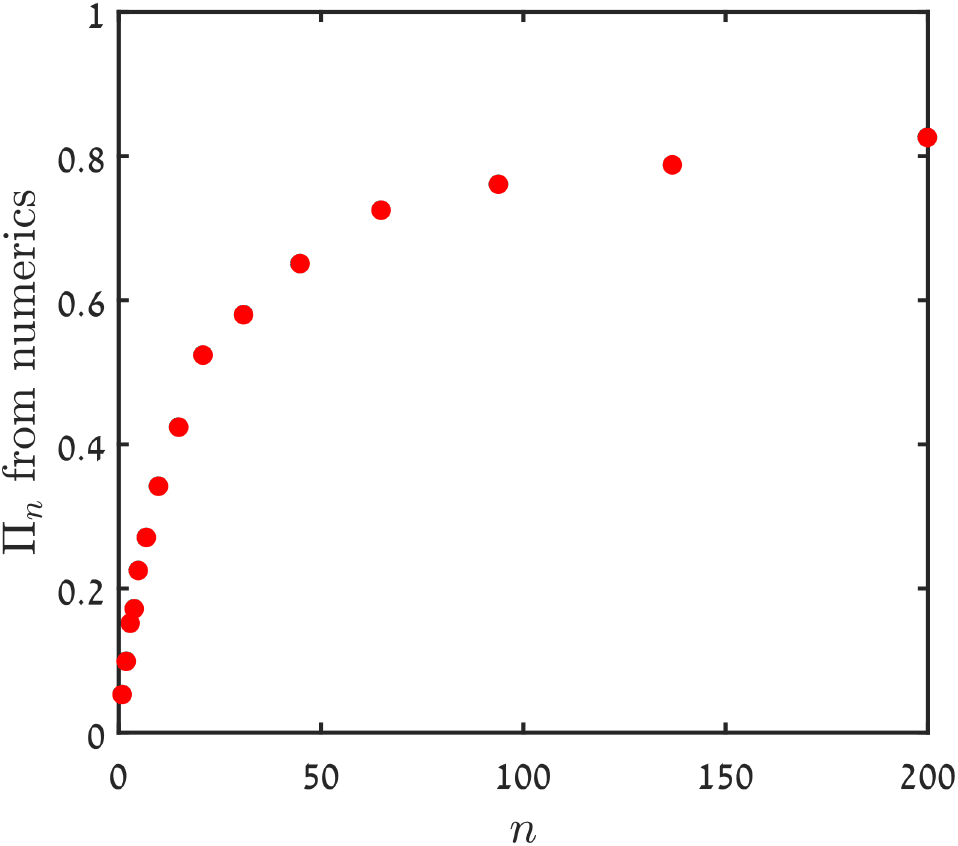
The chance of invasion (*n*_f_ = 1000) for species 1 vs. its initial abundance *n*, averaged over 1000 trials of Monte-Carlo simulations.

**FIG. S8:**
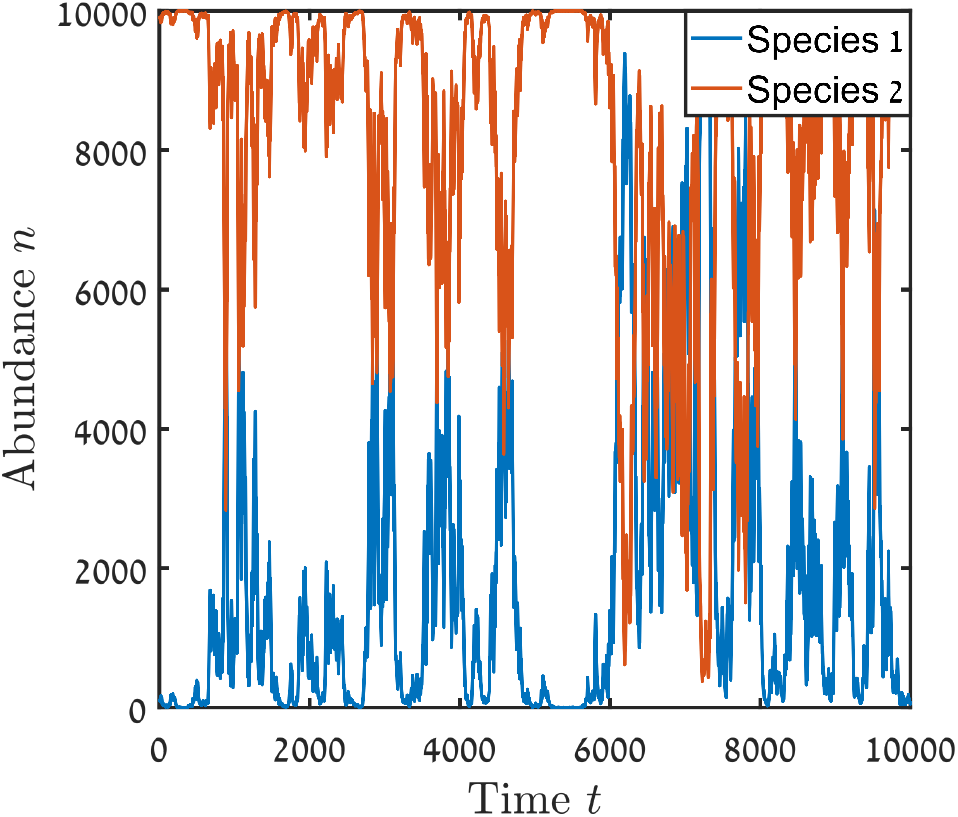
A time series for the abundances of the two species in the forest dynamics model, obtained from the Matlab code presented in subsection F2, between *t* = 0 and *t* = 10000.

These time series were then translated to the logit parameter *z* = log *n*_*t*_/(*N* −*n*_*t*_), in order to find the autocorrelation time of the time series (and, thereby, the dwell time). As explained in [28], the logit parameter is the most suitable for measuring abundance variations at all densities *n*_*t*_/*N*. Note that in all our discussion in the main text we have considered only the invasion regime *n*_*t*_ « *N*, where *z* ≈ ln *n*, but to calculate the autocorrelation time using general time series, the logit variable is to be preferred.

We then plotted the autocorrelation function for the time series of each species (Fig. S9). Assuming that this function decays like exp(−*t/ξ*), we identified *ξ* from a linear regression in a log-linear plot. This step requires some care, since the exponential behavior is sometimes limited to a narrow range of times, but a visual identification of this regime is usually easy. For a given *ξ*, *δ* = 2*ξ* is the dwell time.

**FIG. S9:**
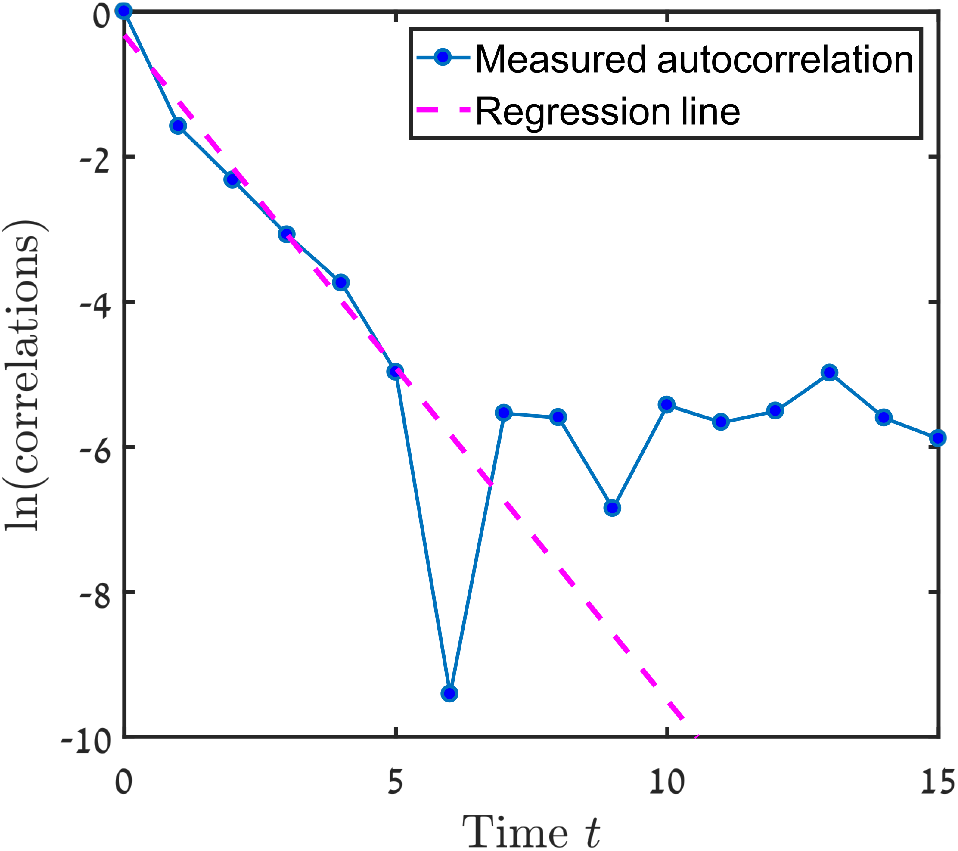
The calculated autocorrelation function is presented on a log-linear plot. Linear regression of the first six points (dashed line) here yields a negative slope of about 0.9, suggesting a dwell time between 2 and 3. We have rounded the dwell time to the higher integer, as the measured autocorrelations in the abundance variations are expected to decay slightly faster than the correlations associated with the actual environmental variations because of the effect of demographic stochasticity.

Once the dwell time *δ* was identified, we calculated the two quantities discussed in Fig. 1 of the main text, namely the mean value 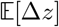 of ∆*z* and the variance Var[∆z] of this quantity for each *n*, and plotted 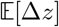 vs. *n* (Fig. S10) and *n* × Var[∆z] vs. *n* (Fig. S11). The code for this is presented in subsection F3. Fitting these plots with straight lines, as explained in the captions of the two figures, we obtained 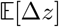, *V*_e_ and *V*_d_. Plugging the values of these parameters into Eq. (2) of the main text we obtained the datasets used to generate Fig. 4 of the main text.

There is some subjectivity involved in the whole process of finding the parameters needed for Eq. (2) from abundance time series, like in the identification of the exponential part of the autocorrelation function or of the regime of *n*-values where *n/N* is small enough to be in the invasion regime but not so small that the initial part where 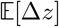 sharply increases with *n* is included (as explained in the caption of Fig. S10). As demonstrated in Figure S12, this subjectivity does not affect the final results too much, and in all the cases we have checked, the final invasibility value has not varied by more than about 20% (and usually much less than that) owing to the differences in measuring the Eq. (2) parameters. Ideally, one should take the mean values of the measured parameters over a few different instances of following the procedure, in order to minimise errors in them.

**FIG. S10:**
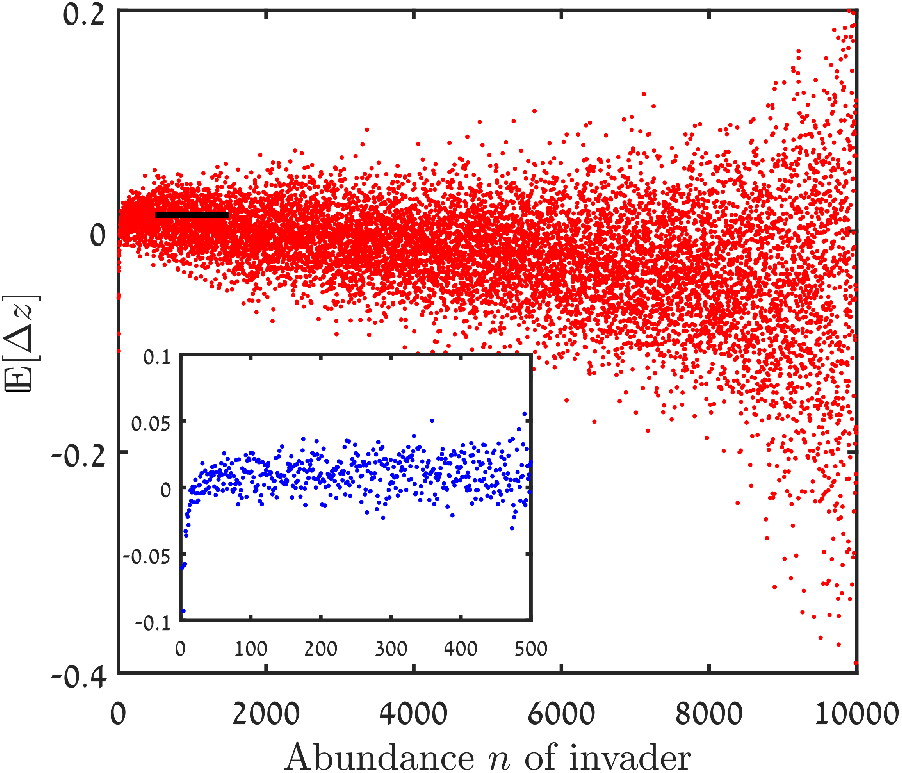
Main panel: 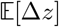 vs. *n*, equivalent to panel (c) of Fig. 1 of the main text. When the small-*n* region of the plot is magnified (inset), a sharp increase in 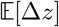 with *n* is observed for very small *n*, due to the effect of demographic stochasticity as explained in the main text. Avoiding these very small values of *n*, we applied linear regression to 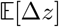 in the small *n/N* regime and used the intercept (thick black line) as the measured value of 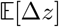.

**FIG. S11:**
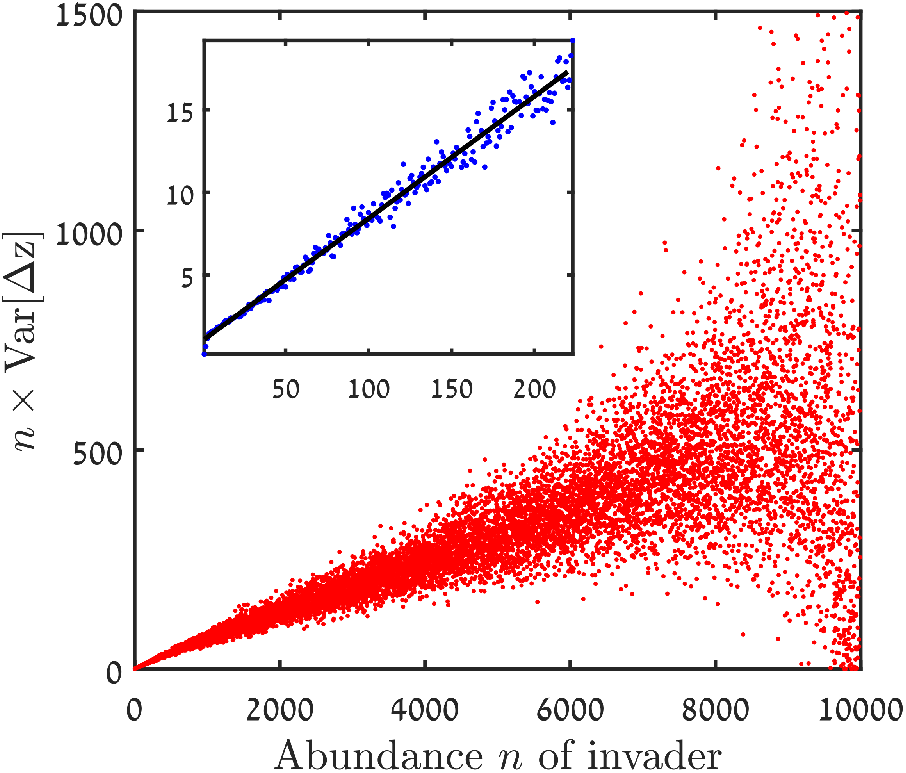
*n* Var[∆z] vs. *n*, equivalent to panel (d) of Fig. 1 of the main text. A linear regression fit at small *n* values (inset, regression line shown in black) yields *V*_e_ as its slope and *V*_d_ as its intercept.

In Figure 4 of the main text we plotted the results for *A* = 0.01 (where *A* is an overall constant that multiplies the recruitment rate time series of each species) and for various values of *f*, *n* and *d* against 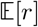 and against our formula (Eq. (2) of the main text). Here (Figure S13) the same figures are presented again. The 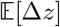, *V*_e_ and *V*_d_ parameters in both the panels were calculated exactly once for each dataset, with 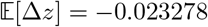 obtained by fitting the regime *n* ∈ [800‥2200] and with *V*_e_ and *V*_d_ found by fitting the regime *n* ∈ [0‥100]. While the resulting agreement obtained between the simulation results and our formula, in Fig. S13(b) (and in Fig. 4(b) of the main text) is not perfect, the deviations are not large, and the predictive ability of our formula is clearly significantly better than that of 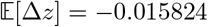 (Fig. S13(a), and Fig. 4(a) of the main text).

**FIG. S12:**
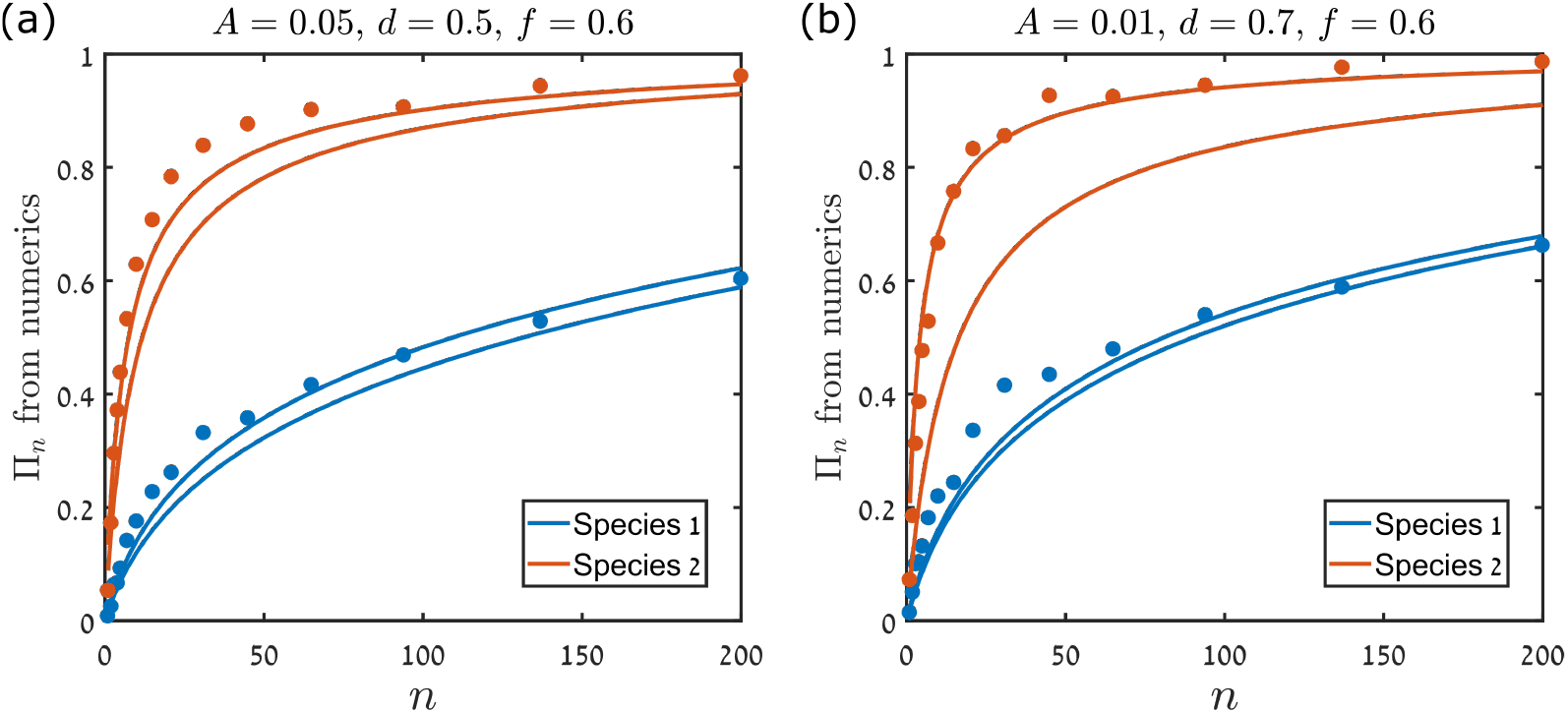
The chance of invasion for the forest dynamics model as measured in numerical simulations (circles) and as predicted by Eq. (2) of the main text (lines). In the title of each subfigure, *A* is an overall constant that multiplies the recruitment rate time series of each species, and *d* and *f* are as defined above. For each set of parameters, different lines correspond to different estimations of the parameters 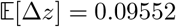, *V*_e_ and *V*_d_, as obtained from two successive applications of the procedure described above (using the code presented in subsections F2 and F3). In panel (a), these estimations yield, for species 1, 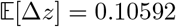, *V*_e_ = 0.29463 and *V*_d_ = 2.281 in one attempt, and 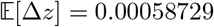, *V*_e_ = 0.2985 and *V*_d_ = 2.118 in the second attempt. For species 2, we obtained 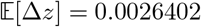, *V*_e_ = 0.2815 and *V*_d_ = 1.9924 in the first attempt and 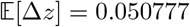, *V*_e_ = 0.31971 and *V*_d_ = 1.3593 in the second attempt. Similarly, in panel (b), for species 1, the first measurement attempt yielded the parameters 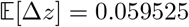, *V*_e_ = 0.13869 and *V*_d_ = 1.4524, while the second attempt yielded 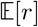, *V*_e_ = 0.14099 and *V*_d_ = 1.4093. For species 2, we found 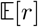, *V*_e_ = 0.15023 and *V*_d_ = 1.5319 in the first attempt and 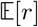, *V*_e_ = 0.16207 and *V*_d_ = 0.44297 in the second attempt.

**FIG. S13:**
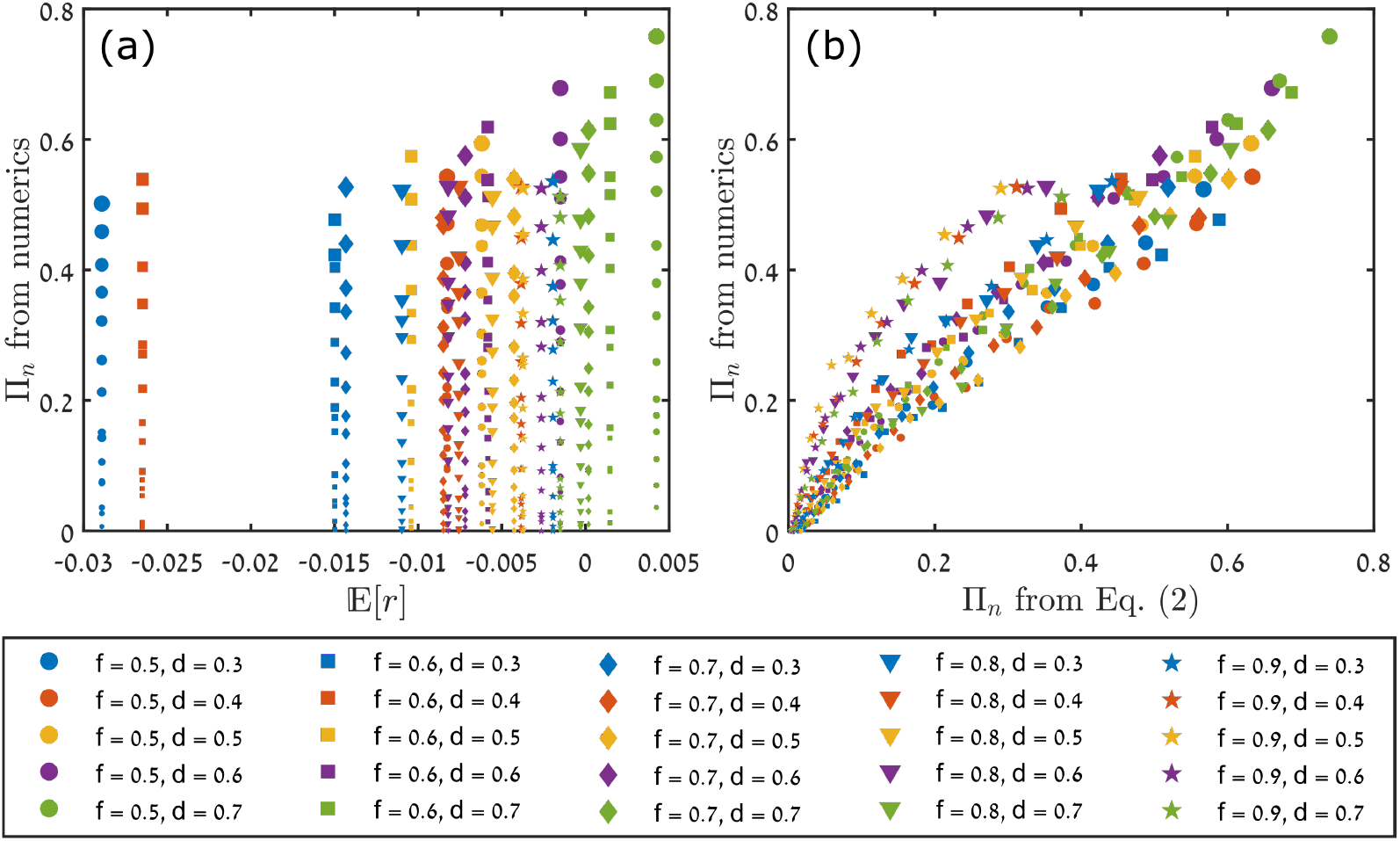
Same as Fig. 4 of the main text.

#### F1. Matlab code for the chance of invasion: Monte-Carlo simulations

Here we provide the Matlab code to find the chance of invasion of a species in the Leslie-Gower model, as explained in section F of this supplement. Explanatory comments in the code are shown in green.

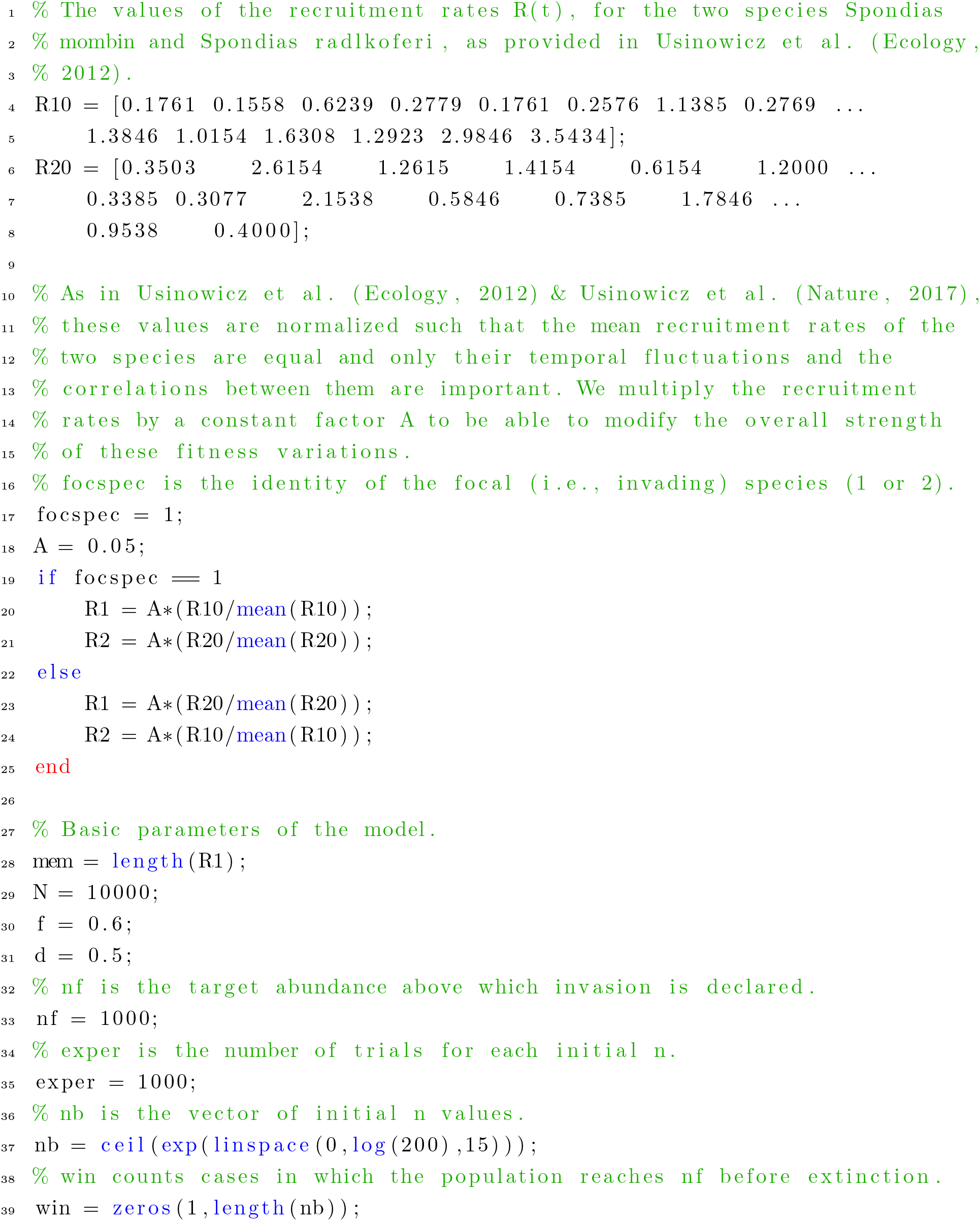
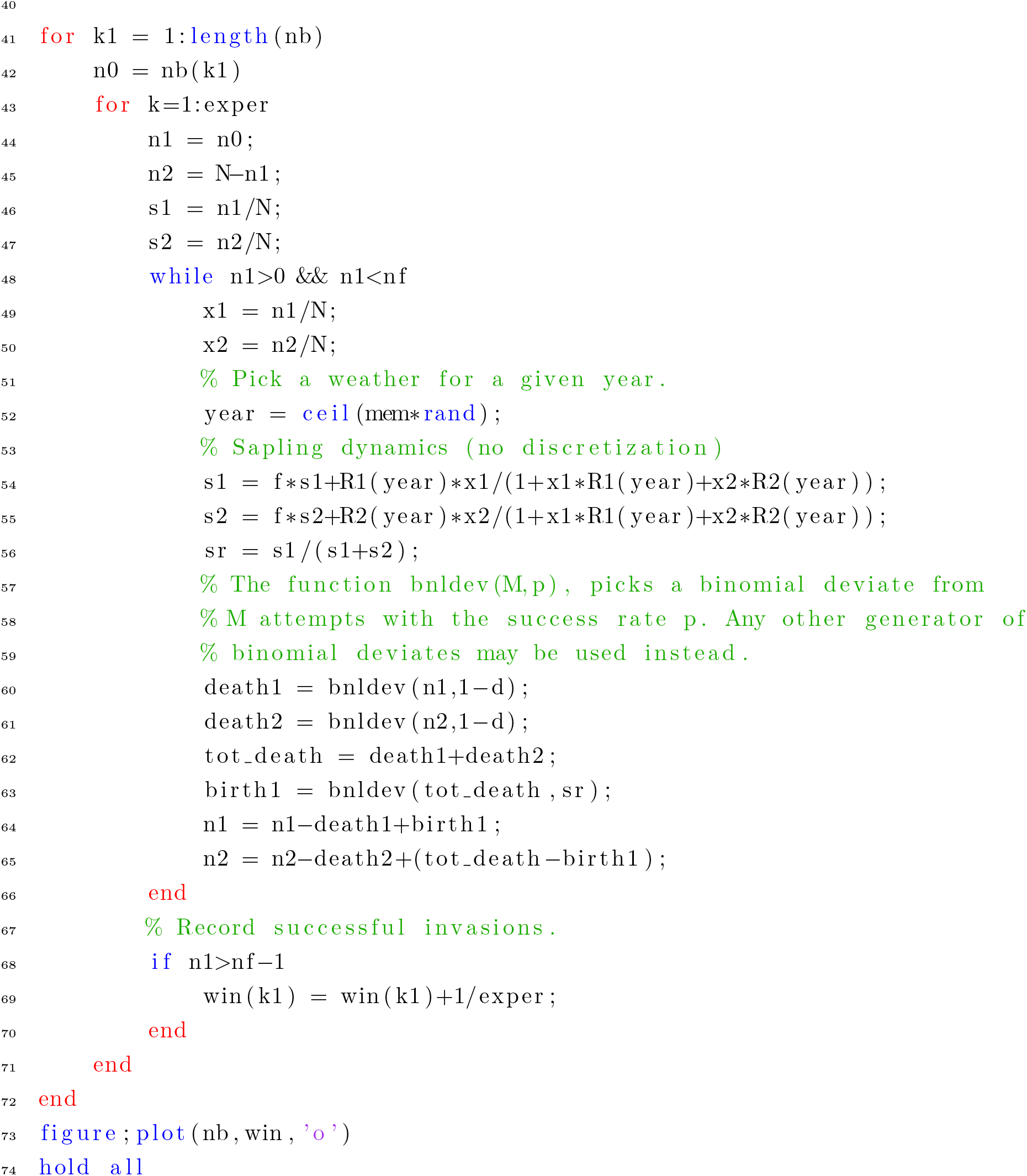

A typical outcome of the above code is shown in Fig. S7, and is repeated here for ease of access.

**FIG. S14:**
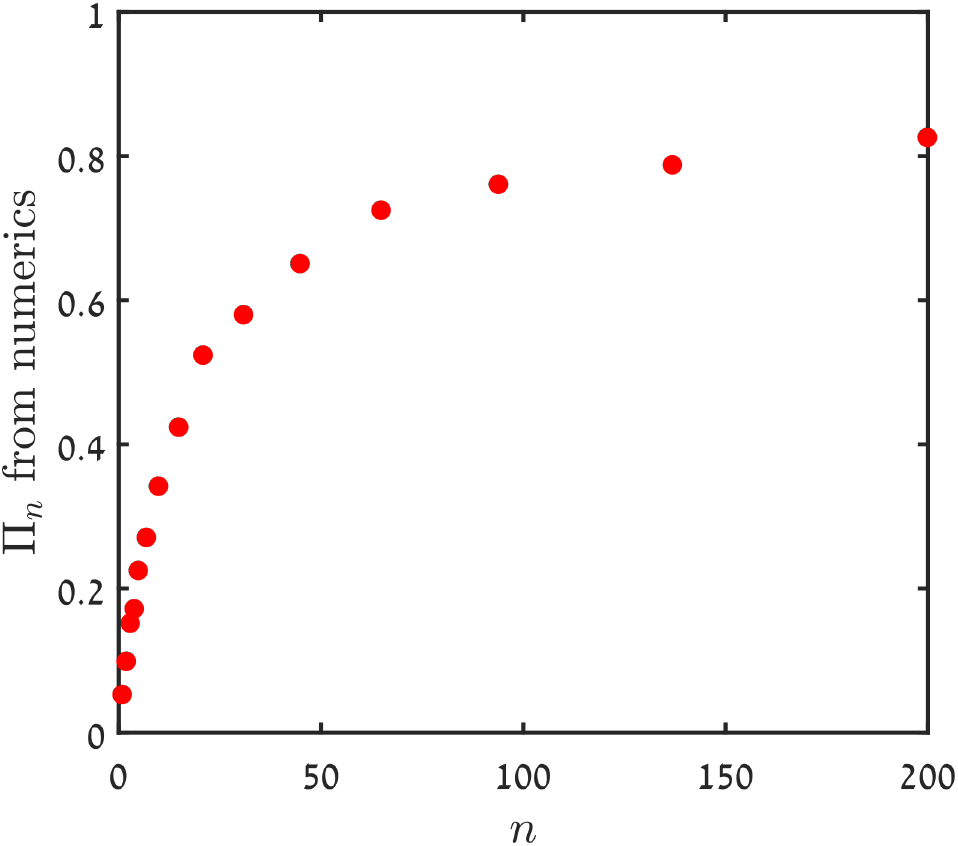
The chance of invasion (*n*_f_ = 1000) for species 1 vs. its initial abundance *n*, averaged over 1000 trials of Monte-Carlo simulations.

#### F2. Matlab code to generate long time series and calculate dwell times

The following Matlab code generates long abundance time series and calculates the autocorrelation function. Explanatory comments are shown in green.

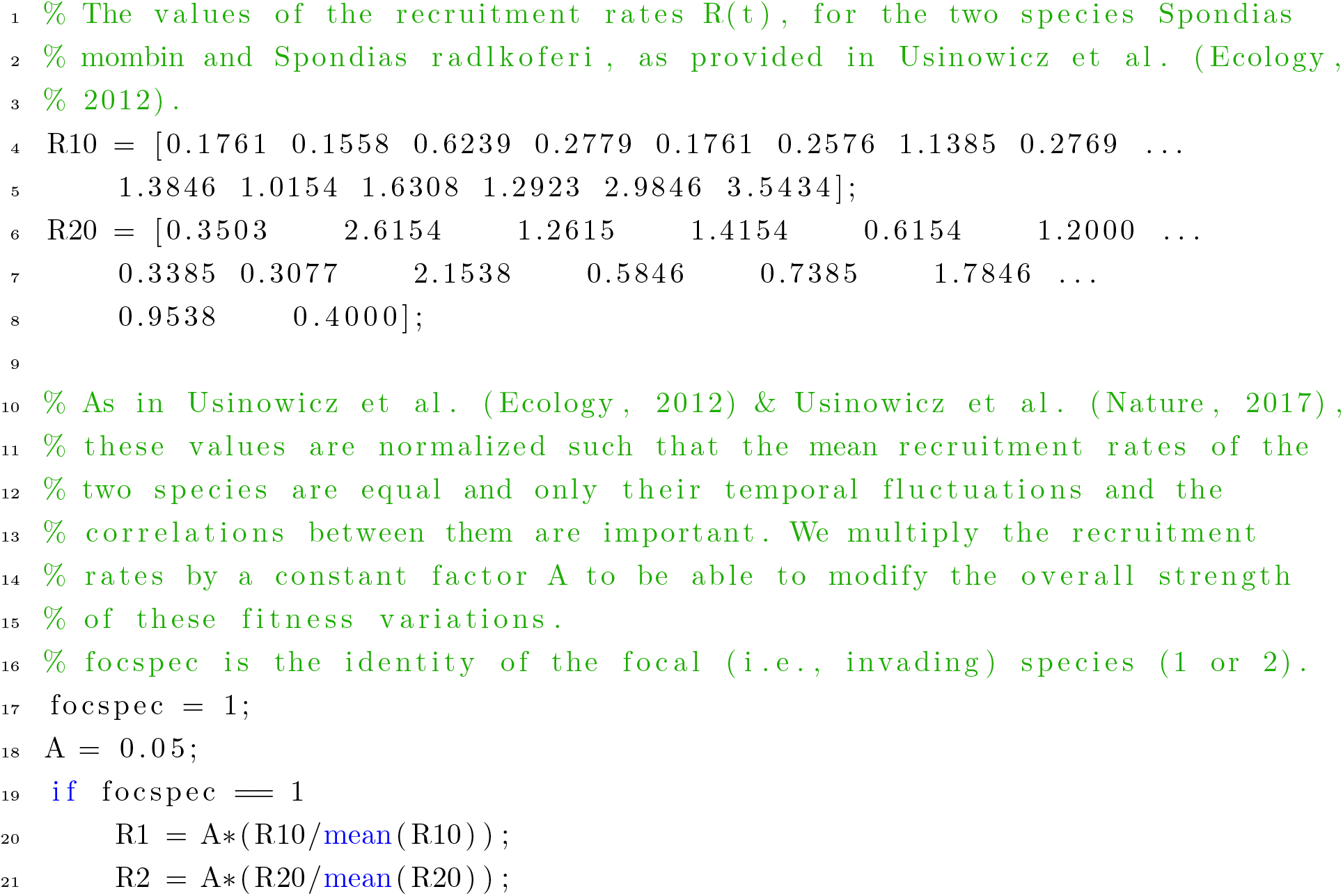
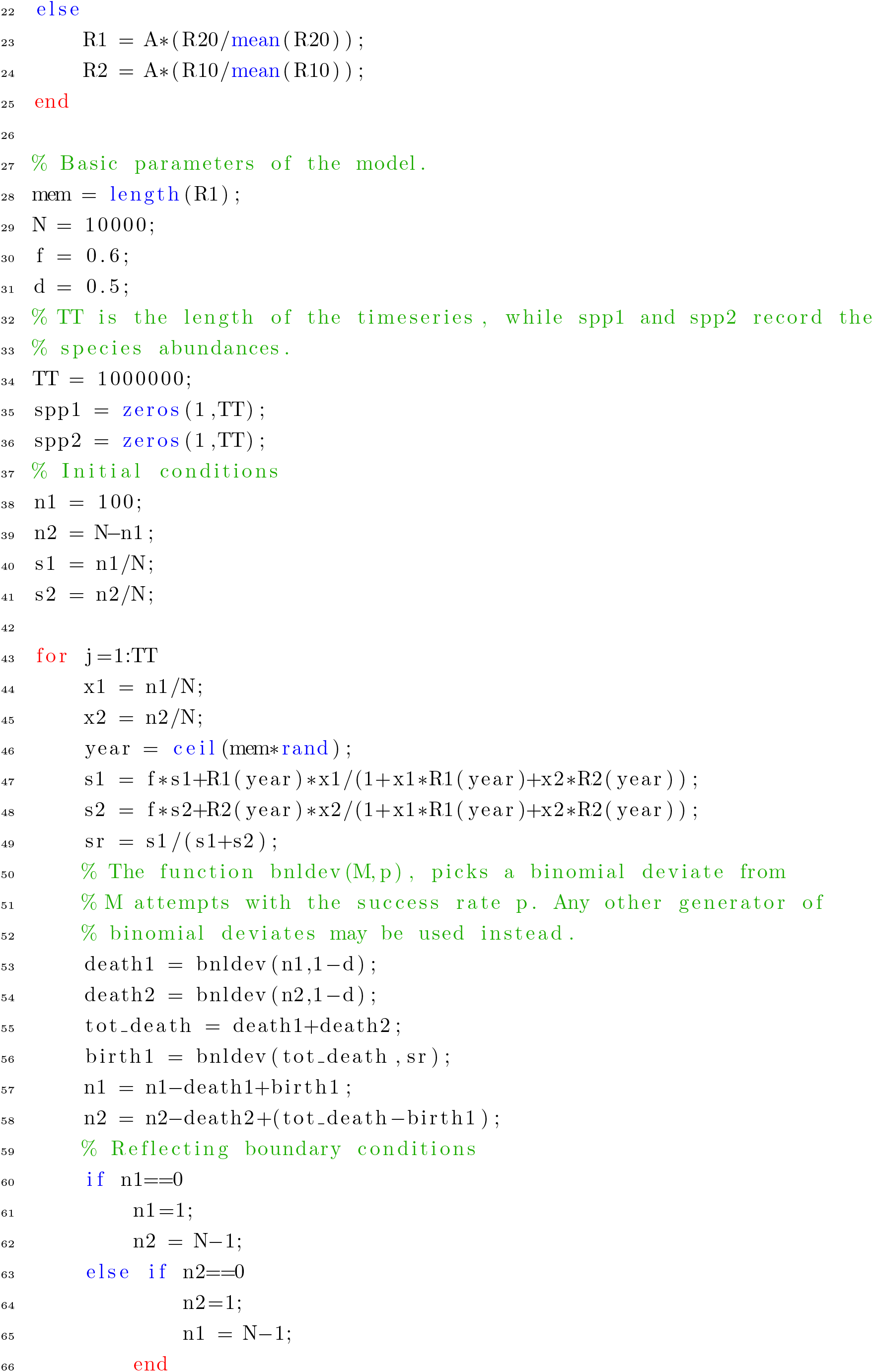
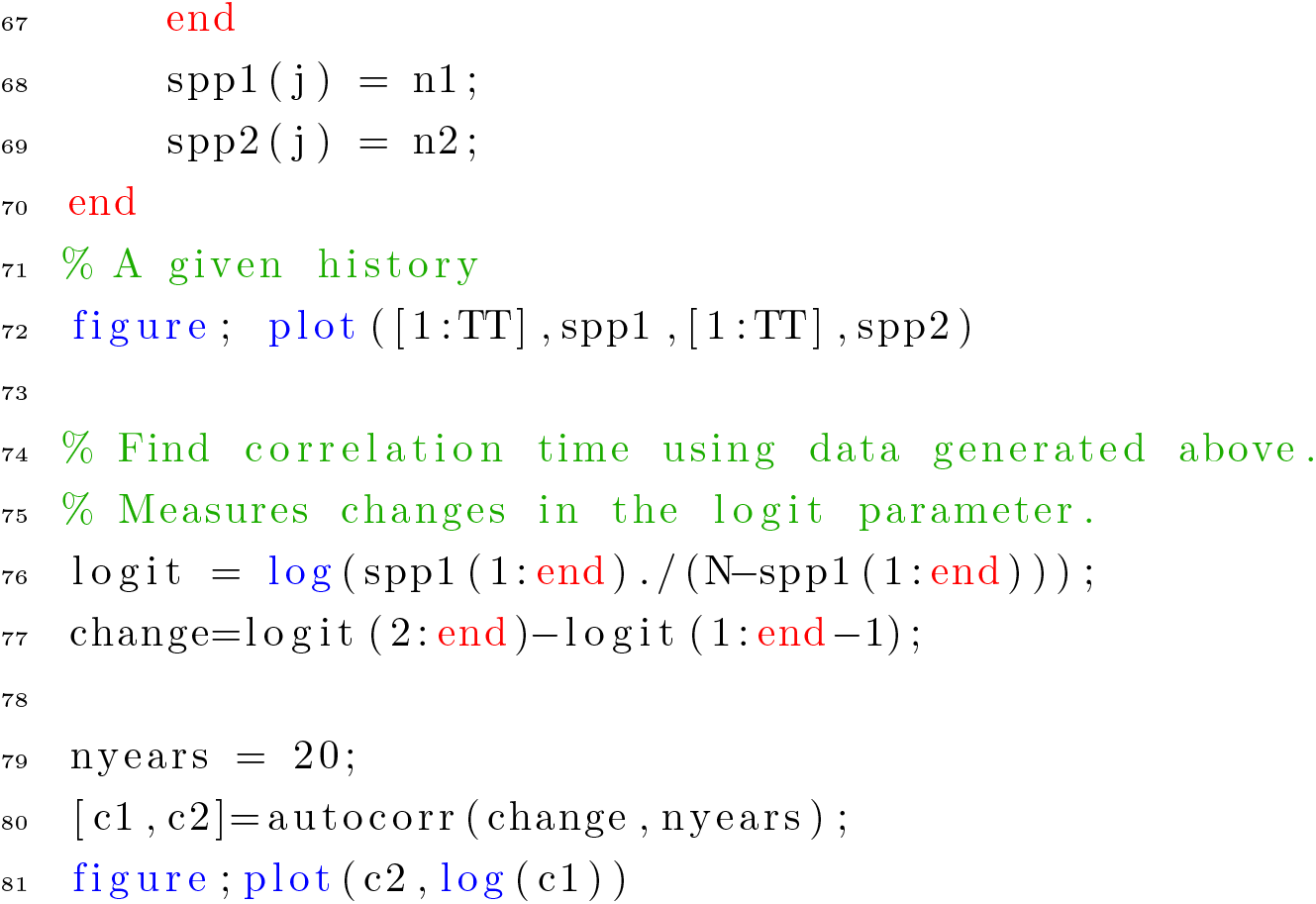

An example of abundance time series for the two species, generated by the above code, is presented in Fig. S8 and is repeated here for ease of access.

**FIG. S15:**
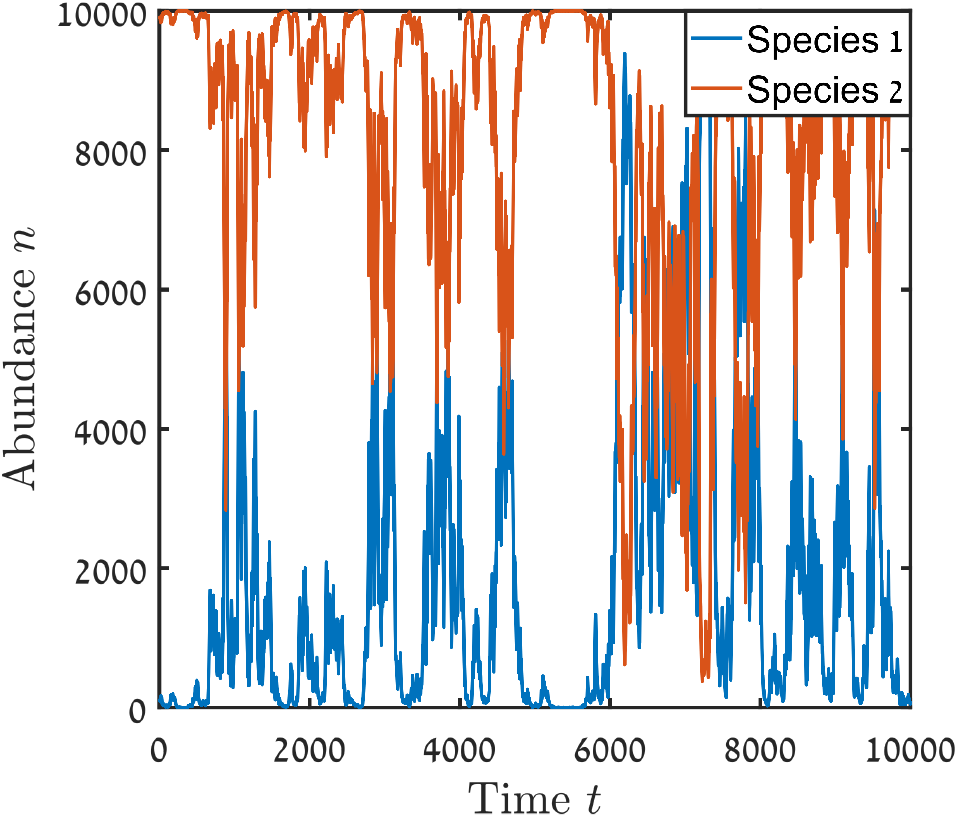
A time series for the abundances of the two species in the forest dynamics model, between *t* = 0 and *t* = 10000.

The autocorrelation function and its fit are shown in Fig. S9, and are also repeated here.

**FIG. S16:**
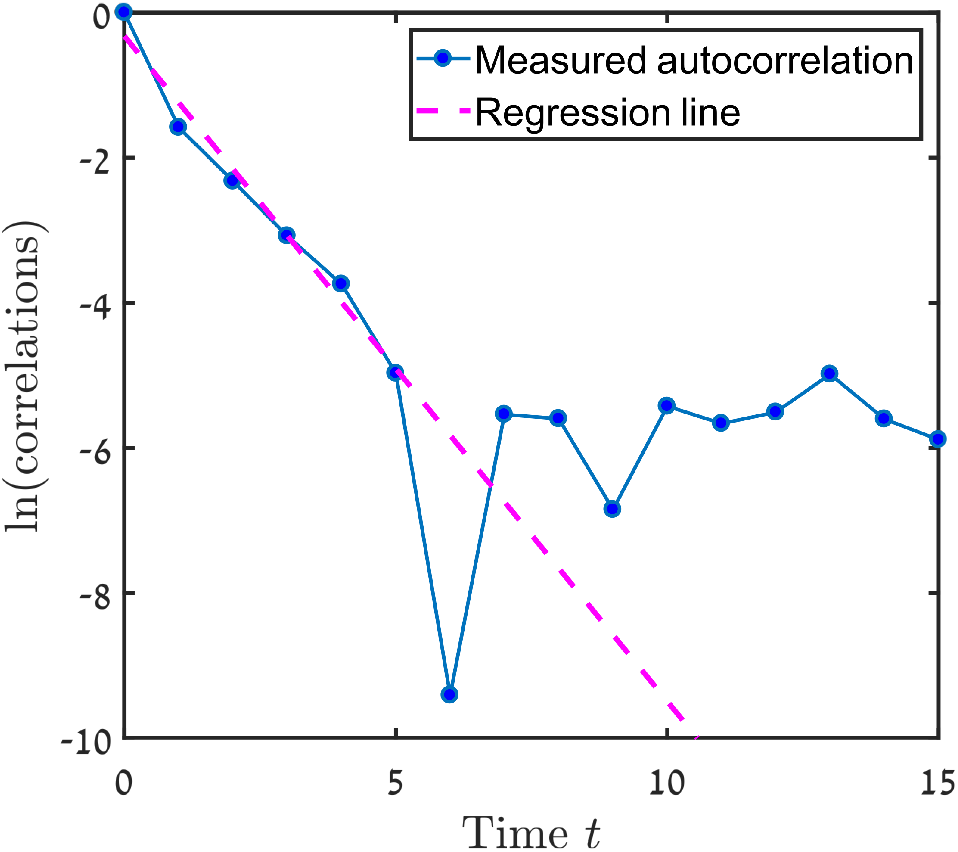
The calculated autocorrelation function is presented on a log-linear plot. Linear regression of the first six points (dashed line) here yields a negative slope of about 0.9, suggesting a dwell time between 2 and 3. We have rounded the dwell time to the higher integer, as the measured autocorrelations in the abundance variations are expected to decay slightly faster than the correlations associated with the actual environmental variations because of the effect of demographic stochasticity.

#### F3. Matlab code to find 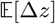, *V*_e_ and *V*_d_

The following Matlab code calculates the mean value 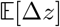 of ∆*z* and its variance for each value of *n*. Again the explanatory comments are shown in green.

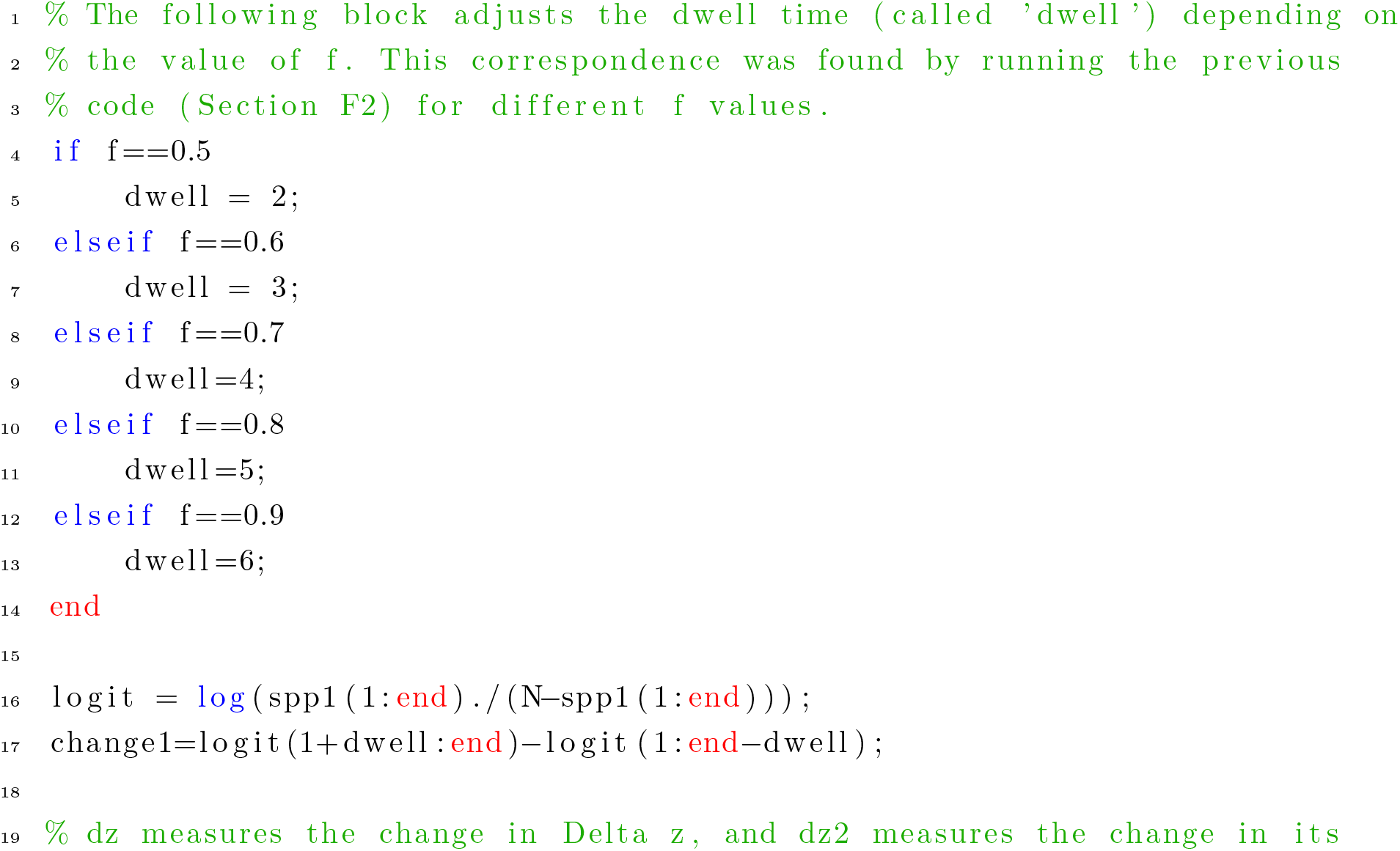
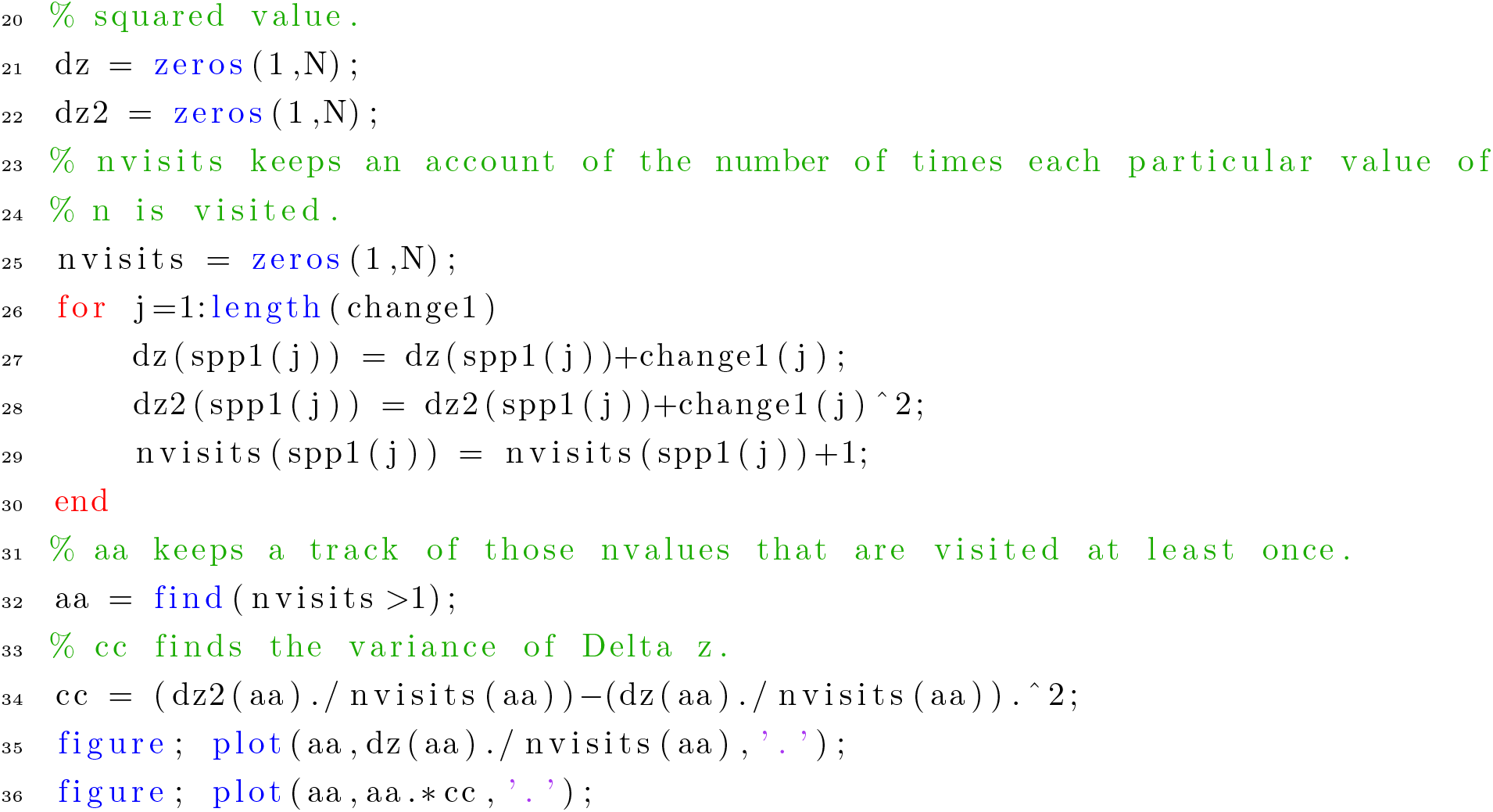

The outcomes of the code above are plots of 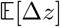 vs. *n* (Figure S10) and *n* × Var[∆z] vs. *n* (Figure S11), which correspond respectively to panels (c) and (d) of Fig. 1 of the main text. For convenience, Figs. S10 and S11 are reproduced here.

**FIG. S17:**
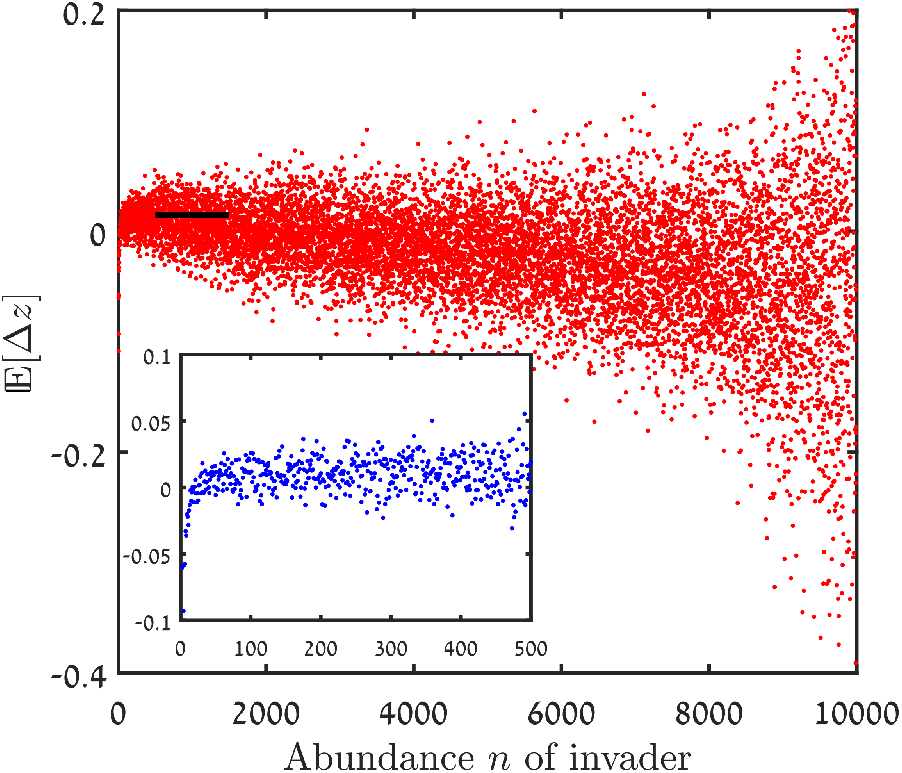
Main panel: 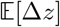 vs. *n*, equivalent to panel (c) of Fig. 1 of the main text. When the small-*n* region of the plot is magnified (inset), a sharp increase in 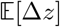 with *n* is observed for very small *n*, due to the effect of demographic stochasticity as explained in the main text. Avoiding these very small values of *n*, we applied linear regression to 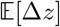 in the small *n/N* regime and used the intercept (thick black line) as the measured value of 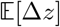.

**FIG. S18:**
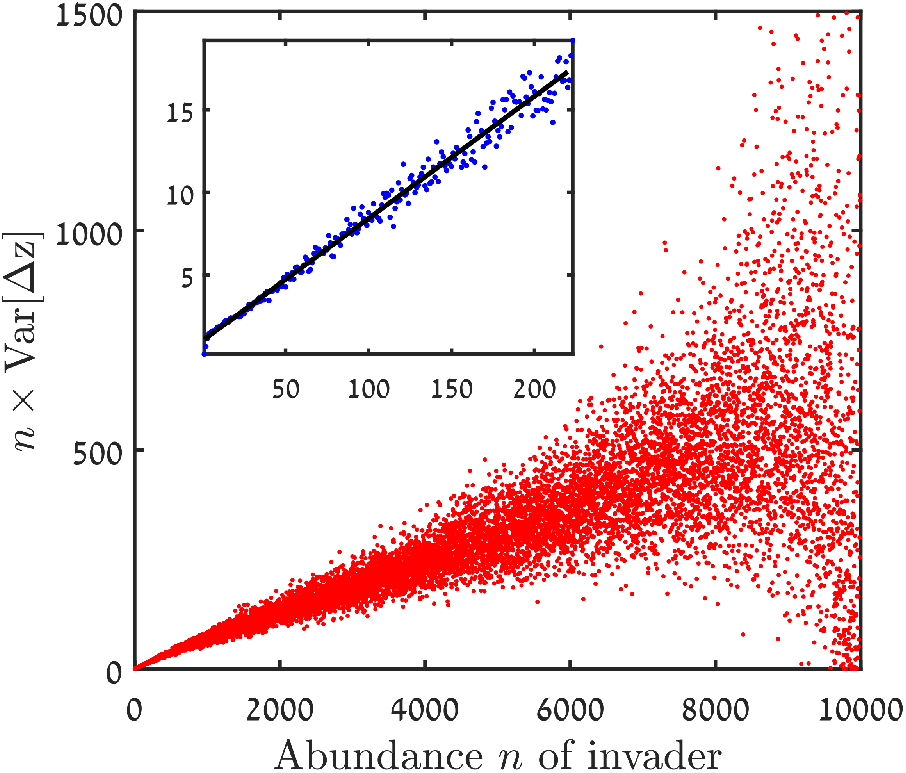
*n* Var[∆z] vs. *n*, equivalent to panel (d) of Fig. 1 of the main text. A linear regression fit at small *n* values (inset, regression line shown in black) yields *V*_e_ as its slope and *V*_d_ as its intercept.

## Notes

### Competing Interest Statement

The authors have declared no competing interest.

